# A Self-Attention Model for Inferring Cooperativity between Regulatory Features

**DOI:** 10.1101/2020.01.31.927996

**Authors:** Fahad Ullah, Asa Ben-Hur

## Abstract

Deep learning has demonstrated its predictive power in modeling complex biological phenomena such as gene expression. The value of these models hinges not only on their accuracy, but also on the ability to extract biologically relevant information from the trained models. While there has been much recent work on developing feature attribution methods that discover the most important features for a given sequence, inferring cooperativity between regulatory elements, which is the hallmark of phenomena such as gene expression, remains an open problem. We present SATORI, a Self-ATtentiOn based model to detect Regulatory element Interactions. Our approach combines convolutional layers with a self-attention mechanism that helps us capture a global view of the landscape of interactions between regulatory elements in a sequence. A comprehensive evaluation demonstrates the ability of SATORI to identify numerous statistically significant TF-TF interactions, many of which have been previously reported. Our method is able to detect higher numbers of experimentally verified TF-TF interactions than existing methods, and has the advantage of not requiring a computationally expensive post-processing step. Finally, SATORI can be used for detection of any type of feature interaction in models that use a similar attention mechanism, and is not limited to the detection of TF-TF interactions.

## INTRODUCTION

High-throughput sequencing techniques are producing an abundance of transcriptome and epigenetic datasets that can be used to generate genome-wide maps of different aspects of regulatory activity. The complexity and magnitude of these data has made deep neural networks an appealing choice for a variety of modeling tasks, including transcription factor (TF) binding prediction (2, 15, 29, 30, 42), chromatin accessibility analysis (3, 4, 18), prediction of chromatin structure and its modifications (19, 34), and identification of RNA-binding sites (24, 41). Besides providing improvement in accuracy over traditional machine learning models, deep learning methods typically require less feature engineering, and can learn directly from sequence and other data. Moreover, deep learning models are able to capture non-linear feature interactions that can help explain the underlying regulatory phenomena.

The discovery that TFs work in tandem to regulate the expression of their targets (39) has sparked the development of a variety of computational methods for predicting cooperativity among TFs and other regulatory proteins by looking at regulatory element co-occurrences (6, 11, 14, 28, 32, 33, 36). In fact, a recent report has demonstrated that TF cooperativity in active enhancers is dominant (31). Despite the demonstrated ability of deep neural networks to extract regulatory signals directly from sequence, there are very few studies that explore cooperativity between regulatory features in genomic data using these methods. Deep Feature Interaction Maps (*DFIM*) uses a network attribution method called DeepLIFT (35) to estimate interactions between regulatory elements, tested for one pair at a time (10). The major drawback of DFIM is that it is computationally expensive: the interactions are inferred in a separate post-processing step and involves recalculation of network gradients. We note that the recent *DeepResolve* method infers feature importance and whether a feature participates in interactions with other features, but does not infer pairs of interacting features explicitly (22).

Recently, neural networks that use the concepts of *attention* and *self-attention* (21, 25) have achieved remarkable success in natural language processing tasks, specifically in machine translation (38). One of the strengths of attention is that it can capture associations between features regardless of the distance between them, addressing a major shortcoming of convolutional and recurrent networks. This is particularly useful for tasks in computational biology where our goal is to identify regulatory elements and their associations/interactions in DNA or RNA sequences. The value of attention for transcription factor binding site prediction was recently demonstrated, motivated by the greater interpretability of the resulting networks (7, 26). However, to the best of our knowledge, attention has not been employed for inferring regulatory interactions between TFs and other regulatory elements.

In this work we propose **SATORI**, a self-attention based deep learning model that captures regulatory element interactions in genomic sequences. The primary components of the architecture of our model are a CNN layer and a multi-head self-attention layer. Optionally, we also incorporate an RNN layer between the two primary layers. The convolutional layer discovers features (motifs) in the input sequences (for caveats and architecture choices that affect this ability see (20)). The self-attention layer then captures potential interactions between those features without the need for explicitly testing all possible combinations of motifs. That enables us to infer a global landscape of interactions in a given genomic dataset without the need for a computationally expensive post-processing step.

We test SATORI on several simulated and real datasets, including data on chromatin accessibility in 164 cell lines in all human promoters and genome-wide chromatin accessibility data across 36 samples in Arabidopsis. To compare our method to DFIM, we incorporated their Feature Interaction Scores (FIS) (10) into our framework. In all our experiments, SATORI and FIS returned highly consistent sets of interactions. We believe this work will assist researchers in improving the interpretability of complex deep learning methods and providing actionable hypotheses for follow up experiments.

## MATERIALS AND METHODS

### Model architecture

We present a self-attention based deep neural network to capture interactions between regulatory features in genomic sequences. Figure 1 shows the model architecture, each aspect of which is described in what follows. The DNA sequences that are the input to the model are represented using one-hot encoding where a sequence of length *L* is transformed into a matrix of size 4 × *L* where each position in the sequence is represented by a column in the matrix with a single non-zero element corresponding to the nucleotide in that position.

**Figure 1.**
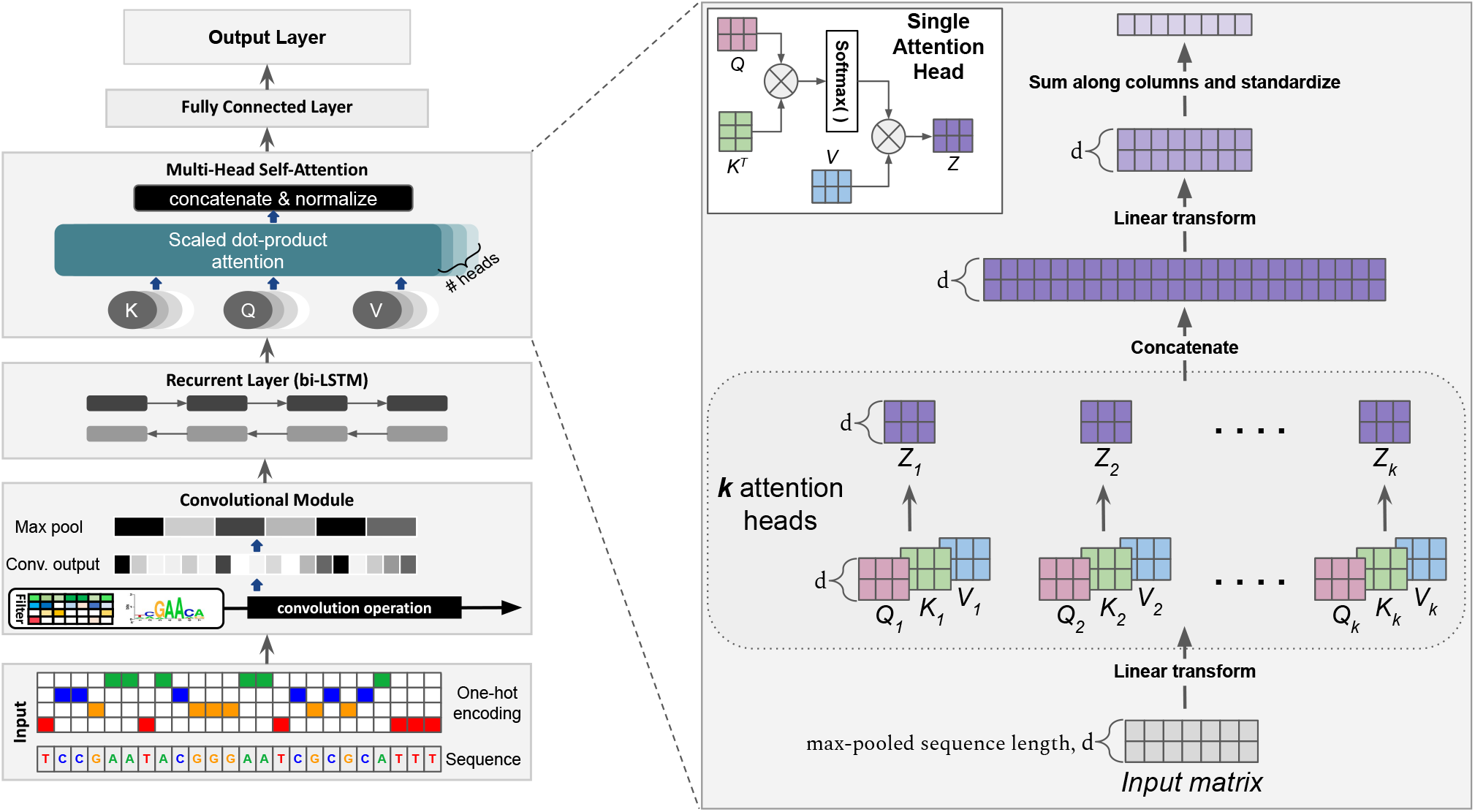
Model architecture: we use a convolutional layer followed by a multi-head self-attention layer; optionally, we add a recurrent layer between the two. The input in both cases is a one-hot encoding of the DNA sequence. The output of the model is either a binary or multi-label prediction. The figure also illustrates the multi-head self-attention layer, details of which can be found in the Supplementary material.

The first component of our model is a one dimensional CNN layer where a set of filters are scanned against the input sequence/matrix. Formally, we can express the result of one dimensional convolution as a matrix *X*′ defined by:

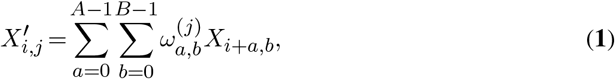

where *X* is the input matrix, *i* is the position at which convolution is performed, *j* is the index of the filter, and *ω*^(*j*)^ is the weight matrix of filter *j*, with size *A × B* where *A* is the length of the filter (window size) and *B* is the number of input channels (four in the case of one-hot encoded input DNA sequences). After the convolution operation, we apply the Rectified Linear Unit activation function (ReLU), which is given by:

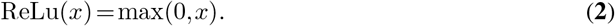

Next, we reduce the output size using max-pooling by taking the maximum value in a window of a pre-determined size, a standard operation in convolutional networks. This reduces the input size for the next layer and also provides invariance to small shifts in the input sequence. We use a relatively small max-pooling window (around six bp) to maintain location information for the subsequent layers, particularly the attention layer which is used to detect interactions.

Following the CNN layer we use an optional RNN layer (see Figure 1). RNNs have an internal state that enables them to collect information across the length of the input sequence. Specifically, we employ a bi-directional RNN with Long Short-Term Memory (LSTM) units (16).

#### Multi-head self-attention

The core component of our network is a multi-head self-attention layer. Attention can model dependencies within the input sequence regardless of their distance (38), a property we leverage to capture TF cooperativity. The key object in self-attention is the *attention matrix*. Consider the input *X* to the attention layer and two linear transformations of it, called the Query *Q*, and Key *K* which are defined by:

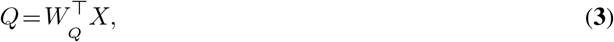

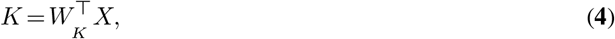

where *W _Q_* and *W _k_* are the corresponding weight matrices for the Query and Key, respectively. The attention matrix *A* is then computed using the following expression:

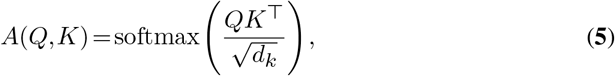

where *d_k_* is the dimension of the Key *K*. The softmax function is applied to each row of the matrix 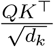, ensuring that the elements of each row sum to 1. The *i*th component of the softmax function applied to a vector **x** is defined by:

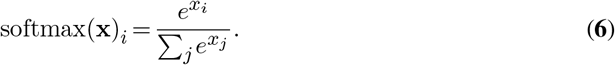

The scaling by 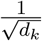 in Equation (**5**) ensures more stable gradients of the softmax function for large sizes of the query/key matrices (38). The attention matrix *A*, defined in Equation (**5**), is a *d × d* matrix where 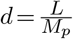*, L* is the length of the input sequence, and *M_p_* is the size of the window used in the max-pooling operation. For every position in the max-pooled output, the corresponding row in the attention matrix summarizes the influence of all other locations on that position. Intuitively, the Query corresponds to a position of interest in the sequence; the dot product between Query and Key compares that position with all other positions in the sequence, summarizing the relevance of all other positions in the corresponding row of the attention matrix. Each row is normalized using the softmax function to normalize the influences gathered from across the sequence. This choice also leads to sparsity of the resulting attention matrix. To generate the output of the attention layer, we first define the Value matrix

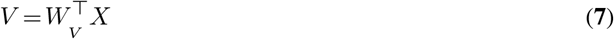

using the associated weight matrix *W_V_*. Finally, we define the output of the attention layer as:

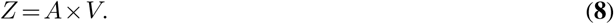

This allows us to generate the output using the parts of the input that we want to focus on—those which exhibit strong inter-dependencies—and ignore irrelevant information.

We then concatenate the individual heads followed by a linear transformation. The final output is then collapsed along the hidden (attention) layer dimensions through addition and normalized by the mean and the standard deviation of the result. Empirically we find that this final step is not only computationally efficient but also leads to better model accuracy in comparison to flattening the attention layer output. For explicit details, please refer to the Supplementary Material. The final fully connected read-out layer outputs the model’s prediction: either binary or multi-label classification, depending on the experiment. For binary classification, we use the standard cross entropy loss function. For multi-label classification, we use the binary cross entropy with logits loss function:

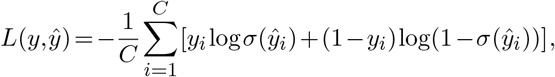

where *y* is the vector of ground truth labels, 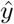 are the network predictions, *C* is the number of classes, and *σ* is the sigmoid function.

### Network training and evaluation

For model selection and optimization, we employ a random search algorithm to tune the network’s hyperparameters. For the convolutional layers we considered filter size, number of filters, and size of window over which pooling is performed. For the multi-head attention layer we tuned the dimensionality of the features generated, and the size of the output of the multi-head attention layer. Details are provided in Table S1 in the Supplementary material. To evaluate the model, we use a simple strategy of splitting the data into 80%, 10%, and 10% for train, test, and validation sets, respectively; we use Area Under the ROC Curve (AUC) to assess model performance. The package was implemented in PyTorch (27) and all the experiments were ran on a Ubuntu workstation with a 12 GB TITAN V GPU.

### Motif extraction

To interpret the deep learning model, we extract sequence motifs from the weight matrices (filters) of the first convolutional layer, similarly to the methodology used in (18). For binary classification problems, we use the positive test set examples that achieve a probability score greater than 0.70. This cutoff was chosen as a good trade-off between the number of qualifying examples and confidence in the prediction. We use all test set examples when dealing with a multi-class or multi-label problems. Next, for each filter we identify regions in the set of sequences that activate the filter with a value greater than half of the filter’s maximum score over all sequences. The resulting substrings are stacked and for each filter, a position weight matrix (PWM) is calculated using the nucleotide frequency and background information. The terms “filter” and the motif (PWM) that describes it are used interchangeably. Sequence logos are generated using the WebLogo tool (9). The PWMs are searched against appropriate TF databases using the TomTom tool (12) with distance metric set to Euclidean. For searching we use the human CISBP (40) and arabiodpsis DAP (23) databases. In the benchmark experiments, we use custom TF databases, details of which are provided in the supplementary material.

### Quantifying feature interactions

In this section we describe the process of inferring filter-filter interactions from the self-attention layer. The attention matrix for each head is calculated using Equation (**5**). Next, we collapse the *k* heads to a single *d × d* matrix by taking the maximum at each position (see Figure 2). This step summarizes the self-attention values from multiple subspaces associated with the corresponding single-heads to a single, attention profile.

**Figure 2.**
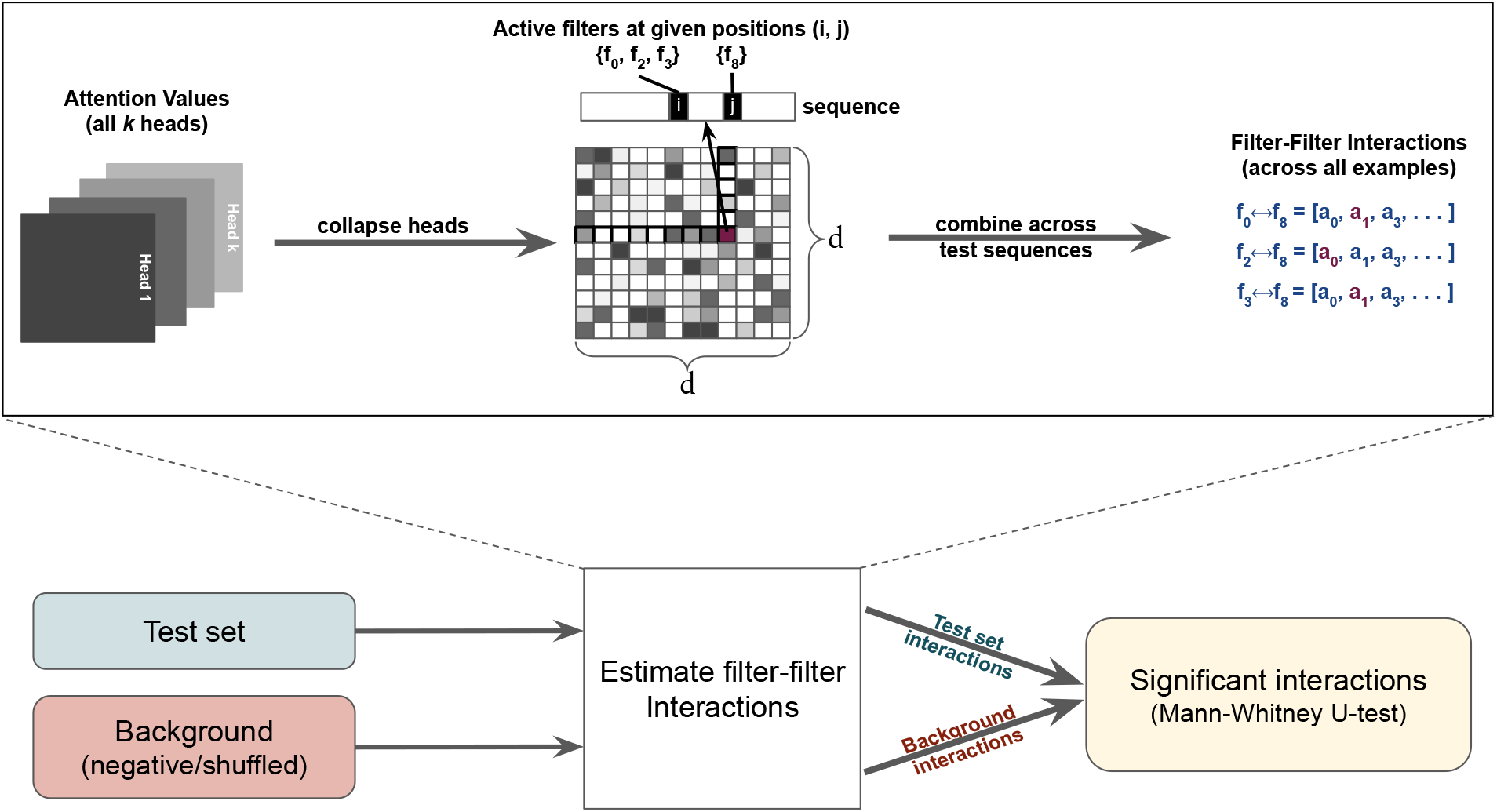
Summary of the process of inferring interactions from self-attention layer values. For a given example, we collapse the attention heads into a single matrix. Next, at each pair of positions, the corresponding active CNN filters are identified and the attention value is assigned to the interacting pair. This is repeated for all examples to generate interaction profiles for all filter-pairs. Finally, we use a background set to test the significance of filter-filter interactions.

The attention matrix provides information about interactions between positions in the sequence. That information is next converted into interactions between filters by retrieving the filters that are active in those positions. Although we perform max-pooling following the convolutional layer, pooling is performed over a very small window (six bp in our experiments) so that sufficient positional information is maintained to accurately detect active filters. The max-pooling operation was useful for reducing the length of the resulting sequences to reduce the computational overhead. Finally, for each identified filter-filter interaction, we generate the attention profile across all testing examples. For a given pair, this profile consists of a vector of its attention values at positions where the corresponding filters were active. An interaction pair is discarded if its maximum attention value is below a certain threshold. By default we used the value 0.10; in the human promoter data we used to increase sensitivity. In the supplementary material we provide the distributions of attention values for all four datasets (see supplementary figure F1).

Filter-filter interactions are then translated to TF interactions by picking the most significant TomTom hits in the appropriate TF database. Filters often learn redundant motifs and, as a consequence, a single TF-TF interaction can be captured by multiple distinct filter-filter interactions. Nevertheless, we observe that for a given TF-TF interaction, the corresponding interacting filters have very similar attention scores (see supplementary figure F8). Note that we might not find significant matches for every CNN filter in the database; it can be expected that our model is capturing interactions of un-characterized regulatory elements. However, in this paper we focus on the interactions between known TFs. We also note that multiple filters can potentially collaborate to contribute to the detection of a single motif pattern. Such cases would be detected as a TF interacting with itself. When reporting TF-TF interactions, the SATORI software ignores such interactions.

To test the statistical significance of motif interactions, we first generate their attention profiles in the background data (described next). Then the non-parametric Mann-Whitney U Test is used to calculate their significance. All the p-values are adjusted for multiple hypothesis testing using Benjamini-Hochberg method (5).

#### Background selection

As mentioned above, to test the statistical significance of regulatory interactions, we need to compare them to a background. We use a biologically relevant background depending on the experiment:

- For binary classification problems, the negative test set is used as the background.
- For multi-label, multi-class or regression problems we generate a background set by shuffling the test

set sequences while preserving their di-nucleotide frequencies. Next, in the shuffled sequences, we randomly embed motifs that are generated based on the CNN filters, interpreted as probability distributions, taking into account the number of times a filter is active above a given threshold in the original test sequences, using the same threshold used for motif extraction.

#### Quantifying interactions using FIS scoring

To infer interactions between motifs using the FIS method, we closely follow the strategy described in (10). Given a test sequence and all of its activated first layer CNN filters, first a source motif is selected. The remaining filters serve as the target motifs for the given source motif. Using Integrated Gradients (37), we then calculate the attribution scores for all the target motifs in the given test sequence. The attribution score determines how important a motif is for the model to accurately predict the test example. Next, the source motif is mutated based on the GC-content of the given sequence and the attribution scores are recalculated for all targets. Finally, to infer interactions, for each source and target pair, the FIS score is calculated as the difference between the attribution scores for each target motif, before and after mutating the source motif. Intuitively, if modifying the source motif affects the attribution of the target motif, this suggests a potential interaction, and the magnitude of the change in attribution scores is used to quantify this potential. We compute FIS scores for all unique pairs of source and target motifs across all test sequences and identify statistically significant interactions using the the same approach used in SATORI (see Figure 2).

#### Selecting test examples

To quantify interactions using SATORI or the FIS-based approach, we use the high-confidence predictions of the model. For binary classification, we pick all positive examples that are assigned prediction confidence above a specified threshold. We use a threshold of *p* = 0.70 in our experiments. For the background examples, we pick all the negative test examples that score below 1 − *p*. In case of the multi-label classification problem, we pick our test examples based on the precision of the model’s prediction probabilities: for a test example to qualify, the precision value—calculated using the given labels and their model assigned probabilities—must be above a specified threshold (default precision threshold = 0.50). We note that for FIS scoring of multi-label classification problems, we only use the attribution values of the true positive predictions. These values are summed and used in calculating the final FIS score.

### Data collection and processing

We use four datasets to evaluate the ability of SATORI to capture interactions between regulatory elements. The datasets are summarized in Table 1, and specific details are provided in the Methods section of the supplementary material.

**Table 1.**
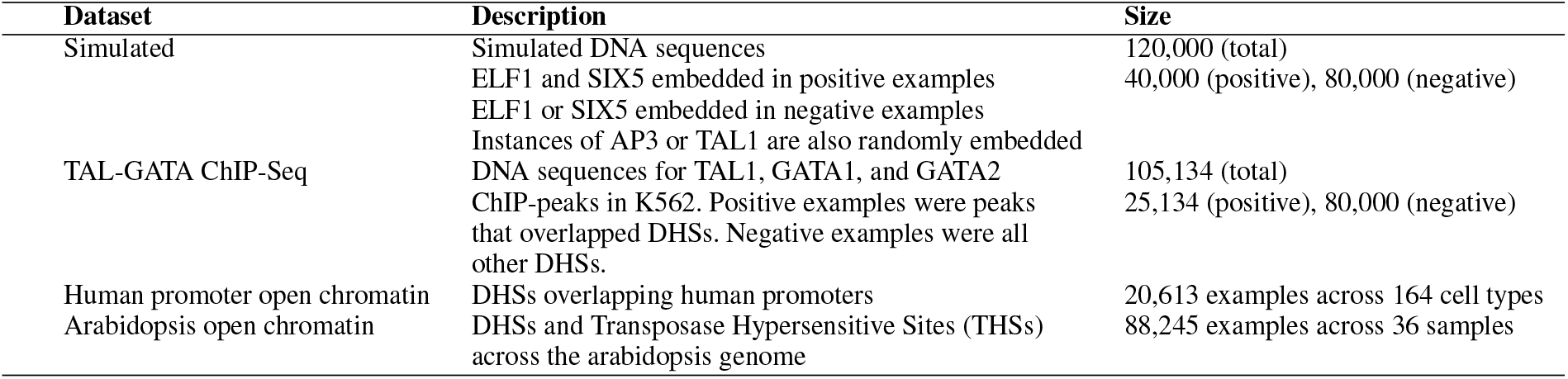
Dataset summary. The first two datasets are binary classification problems whereas the last two experiments are multi-label problems.

## RESULTS AND DISCUSSION

### Benchmark 1: embedded motif interactions in simulated genomic sequences

In this experiment we used SATORI to test if it can recover interactions embedded in simulated DNA sequences. The dataset was created similarly to the simulated dataset of Greenside et al. (10): 120,000 random DNA sequences were generated; in 40,000 sequences we embedded motifs of the transcription factors SIX5 and ELF1. Motifs from both TFs were present in each sequence, simulating an interaction. These sequences were labelled as positive examples. In the rest sequences we embedded only one of the two motifs in each sequence. In addition, motifs occurrences of TAL1 and AP3 were embedded at random across the whole dataset.

Not surprisingly, both variants of our deep learning architectures achieved perfect classification accuracy on the test set for this data. We then analyzed the attention layer weights and inferred statistically significant motif interactions and found that all the significant interactions returned by our model involve SIX5 and ELF1 as expected. These interactions are summarized in Supplementary Table S2.

### Benchmark 2: Inferring TAL-GATA motif interactions from ChIP-Seq data

The TFs TAL1 and GATA1 have been reported to interact: GATA1 requires a prior or simultaneous binding of TAL1 before it can bind DNA (17). To investigate these interactions in this experiment, which follows a similar experiment performed by the authors of DFIM, we formulated a binary classification problem where the positive set consisted of sequences of the TAL1, GATA1, and GATA2 ChIP-Seq peaks that overlapped regions of open chromatin (DHSs) in the human K562 cell-line by at least 200 bp. For the negative set, sequences of all other K562 DHSs that didn’t overlap any of the ChIP-Seq peaks were used. This experiment serves as another benchmark for our model, and was also used by Greenside et al. to test their model’s ability uncover interactions between TAL1 and GATA1/GATA2 (10). Further details regarding the dataset are provided in the supplementary material.

We trained both variants of our model on this dataset; in this harder dataset the variant with an RNN layer performed much better with an AUC of 0.94 on the test set compared to 0.85 for the model without an RNN. The authors of DFIM achieved similar accuracy to our model that uses an RNN using five layers of convolution. Please note that we do not seek to demonstrate better model accuracy — our focus is on model interpretability, which we seek to achieve without compromising on accuracy. For that reason, we chose to present results using the architecture that uses the additional RNN layer. Our experiments with both architectures have yielded very similar results when it comes to detecting interactions (see Figures F6 and F7 in the Supplementary Data). Using SATORI we recovered multiple significant filter-filter interactions that mapped to TAL1 and GATA motifs with highly significant p-values (see supplementary table S3), demonstrating the ability of our model to recover biologically relevant interactions between TFs.

Since this dataset uses ChIP-Seq peaks of TAL and GATA transcription factors that occur in regions of open chromatin, other TF-TF interactions might present in these regions. Therefore, we also let SATORI search for interactions among all known human transcription factors. As expected, we found numerous other interactions. Table 2 summarizes the 15 most frequent TF family-family interactions, consisting predominantly of interactions between members of the C2H2 ZF, Homeodomain, CxxC, and GATA families (see Supplementary Table S4 for the list of individual TF interactions). An interesting observation is that the interactions between the GATA and bHLH (TAL1) families are not the most frequent, despite the fact that the model used their ChIP-Seq peaks. This is likely because of the differences in size of these TF families: the C2H2 ZF and Homeodomain families are the largest TF families in humans, whereas bHLH and particularly GATA, are much smaller in size.

**Table 2.** The most frequent interacting families of human transcription factors in the TAL-GATA ChIP-peaks in human K562 cell-line. All interactions are significant with adjusted p-value < 0.05.

| TF Family Interaction | Frequency | Percent of total interactions |
| --- | --- | --- |
| C2H2 ZF ⟷ CxxC | 170 | 14.36% |
| Homeodomain ⟷ C2H2 ZF | 121 | 10.22% |
| C2H2 ZF ⟷ C2H2 ZF | 90 | 7.60% |
| GATA ⟷ C2H2 ZF | 79 | 6.67% |
| Homeodomain ⟷ CxxC | 72 | 6.08% |
| C2H2 ZF ⟷ Sox | 62 | 5.24% |
| C2H2 ZF ⟷ bHLH | 53 | 4.48% |
| GATA ⟷ CxxC | 51 | 4.31% |
| Sox ⟷ CxxC | 39 | 3.29% |
| C2H2 ZF ⟷ THAP finger | 21 | 1.77% |
| CxxC ⟷ bHLH | 19 | 1.60% |
| Nuclear receptor ⟷ C2H2 ZF | 19 | 1.60% |
| **GATA ⟷ bHLH** | 18 | 1.27% |
| Homeodomain ⟷ bHLH | 17 | 1.44% |
| SAND ⟷ C2H2 ZF | 17 | 1.44% |

Because of the similarity between some TF motifs, the matching between filters and motifs is not without errors. In this second experiment GATA was predicted to interact with TCF15, which has a motif that closely resembles that of TAL1. In fact, both of them belong to the same bHLH family. Figure F2 in the Supplement shows the similarity between these motifs.

### The TF interaction landscape across human promoters

In this experiment we investigated regulatory interactions between TFs in all human promoter regions using DNase z I hypersentivity data (DHSs) across 164 immortalized cell lines and tissues. This experiment was based on the work of Kelley et al. (18) which used convolutional networks to predict chromatin accessibility from sequence information alone across the entire human genome. We chose to focus on a subset of the data that consists of DHSs in human promoters to determine interactions relevant to the regulation of gene expression. The labels in this data represent presence/absence of a given DHS across each of the 164 cell lines, and is a multi-label classification problem. Additional details are found in the supplementary material. We trained both network variants (see Figure 1) and observed that with the optional RNN layer, the network performed better in terms AUC scores (see Supplementary Figure F3). SATORI results are presented for this architecture variant.

The trained network yielded filters that matched 93 TFs with known motifs (counting only filters that had information content greater than 3.0). Among those 93 TFs, our model identified 123 unique pairs of motifs that interact. The 15 most frequent interactions are shown in Figure 3(a). For the complete list, refer to Supplementary Table S5. We also looked at the distribution of the distances between interacting motifs, and observed that, as expected, interactions tend to occur in close proximity with a median distance of interaction of 150 bp (see Figure 3(b)). Overall, the Homeodomain, C2H2 ZF, Sox, and CxxC families were the most frequent families of interacting TFs (see supplementary figure F5). Finally, it is worth mentioning that for four interactions out of the total of 123, we found evidence in the TRRUST database (13) which lists 16 interactions among the 150 TFs. This overlap is statistically significant with a p-value of 5.98 × 10^−4^ using the hypergeometric test. We also found support for nine out of the 123 interactions in the HIPPIE database of human protein-protein interactions (1), which had 56 total interactions among the 150 TFs. This overlap is statistically significant with a p-value of 7.72 × 10^−22^ using the hypergeometric test. Table S6 in the Supplementary Material provides the interactions supported by these two databases. We note that cooperativity between TFs does not necessarily occur through direct physical interactions. Other modes of cooperativity include competitive binding and remodeling of chromatin (8). In this work we only analyzed interactions between motifs of known TFs. Not all filters can be mapped to characterized regulatory proteins. In this dataset, TomTom returned no significant matches for 80 out of the 200 CNN filters with motif information content greater than 3.0. The interactions of these filters require further investigation to discover the regulatory molecules associated with them.

**Figure 3.**
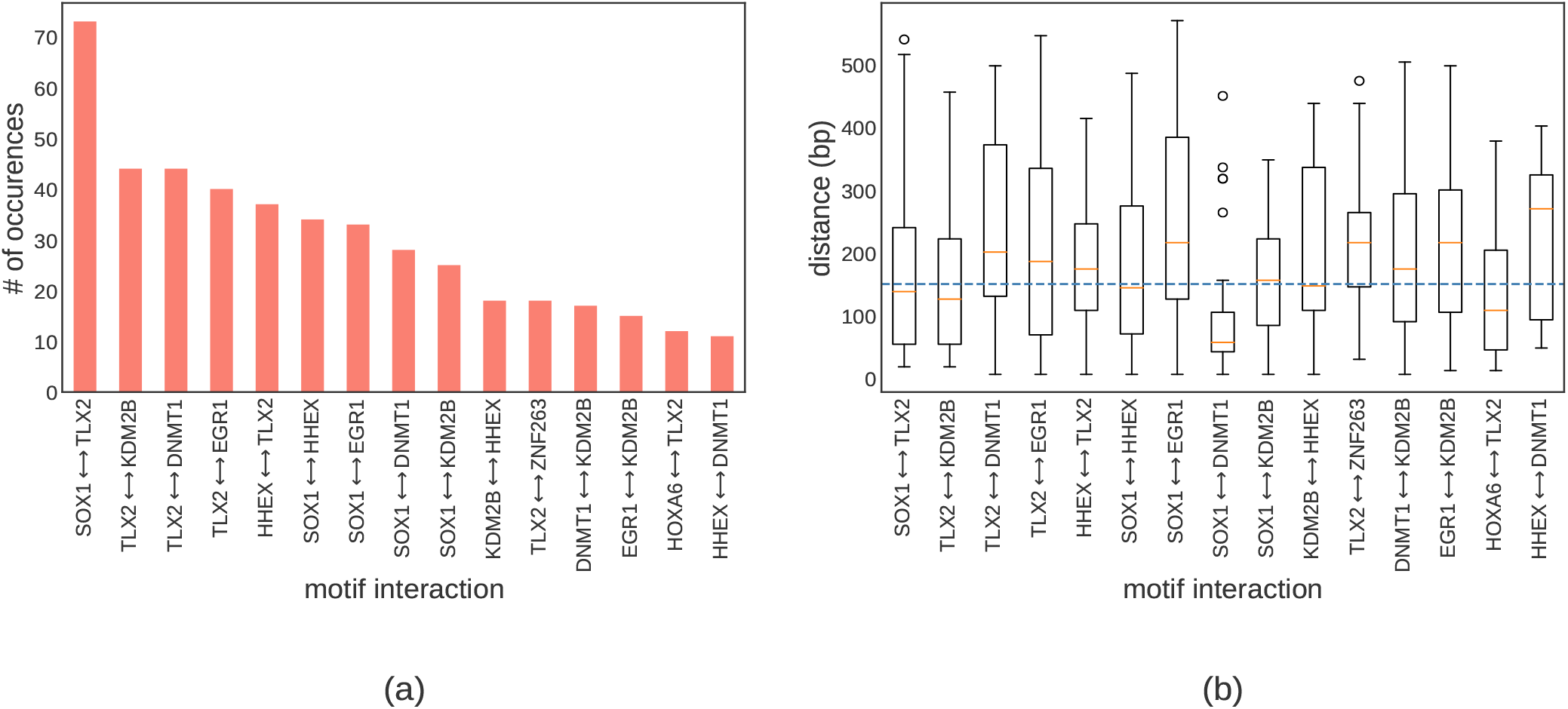
The most frequent TF interactions in human promoters (a). The distribution of TF-TF interaction distances for the most frequent interactions (b). The dotted blue line represents the median distance across all significant interactions.

As mentioned above, the motif matching results returned by TomTom are noisy and imperfect. For example, some of the statistically significant matches are clearly incorrect, as shown in Supplementary Figure F4. However, this is not a shortcoming of our model, but rather a limitation of the interpretation of filter-filter interactions.

### Genome-wide regulatory interactions in arabidopsis

In the next experiment we evaluated the ability of SATORI to detect interactions on a genome-wide scale. For this task we chose to focus on regions of accessible chromatin in arabidopsis in a manner similar to our experiment in human promoter regions. More specifically, we predict chromatin accessibility from sequence across 36 arabidopsis samples from recently published arabidopsis DNase I-Seq and ATAC-Seq studies (GEO accession numbers provided in the Supplementary Methods section). Like the previous dataset, this too is a multi-label prediction problem, where the labels indicates whether a given region has a peak in each of the 36 samples of DNase I-Seq and ATAC-Seq. We trained both deep learning architectures and as in the other datasets, the network that included an RNN layer performed better in terms of median AUC across samples (0.86 compared to 0.85).

In the next step, we investigated genome-wide regulatory interactions in those regions of open-chromatin. The trained network yielded 189 filters with information content above 3.0, and we obtained 100 unique matches for those filters in the DAP-Seq arabidopsis TF database (23). Among these 100 TFs, our model identified interactions between 687 pairs of of motifs involving diverse plant transcription factors (see Figure 4(a)). G2like, MYB, C2C2dof, and AP2 were the most frequently represented TF families in those interactions. Similarly to our findings in human, plant TF interactions tend to occur in relative proximity (median distance = 126 bp) as shown in Figure 4(b). Arabidopsis does not have a database of known interactions between TFs, so our results could not be validated. Targeted experimental validation of these predictions can thus significantly enrich our knowledge of the combinatorial regulation of gene expression in plants.

**Figure 4.**
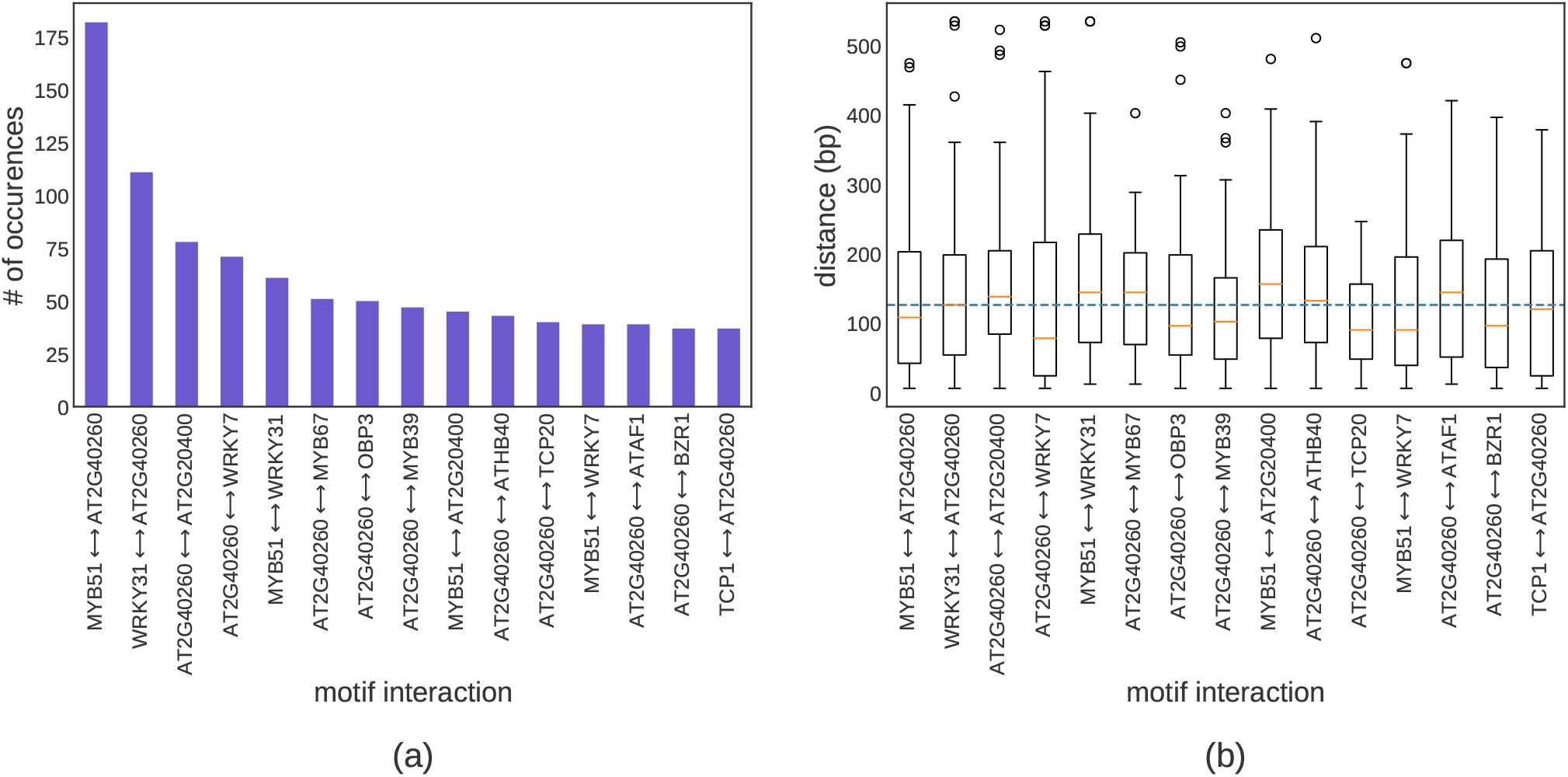
The regulatory interaction landscape in accessible chromatin in the arabipdosis genome. The most frequently interacting families of plant transcription factors (a). For the most frequent interactions, the box and whiskers plot depicts the distribution of interaction distances (b).

### Comparison: SATORI and FIS-based interactions

To compare our model to DFIM (10), we incorporated its FIS scoring method as a feature in our framework and tested it on the three real-world datasets. A key observation is that among the top scoring interactions detected by the two methods there is very high overlap: In the TAL-GATA dataset twelve out of the top fifteen interactions detected by FIS scoring were also reported by SATORI; for the human promoter dataset 14 out of the top 15 FIS predictions were detected by SATORI; finally, for the arabidopsis genome-wide dataset, all the top 15 FIS predictions were reported by SATORI. At the TF family level, we observed perfect agreement in the top ten predictions in all three datasets. These results are summarized in Figure 5; a summary of the number of statistically significant interactions detected by the two methods is provided in supplementary Table S7. The agreement on the top predictions suggests their high likelihood of being biologically relevant, and make them promising candidates for experimental validation.

**Figure 5.**
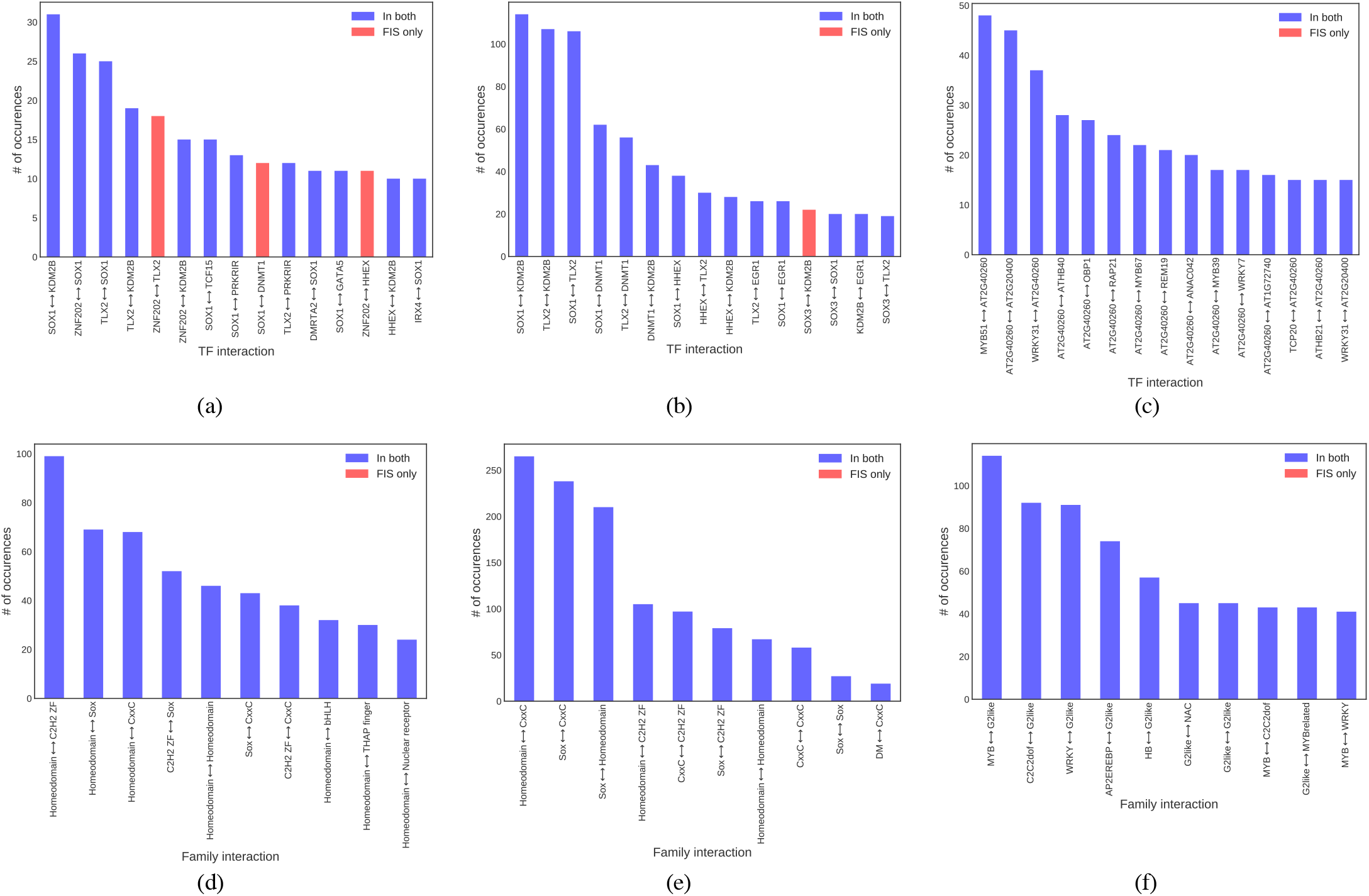
Common interactions in the top predictions of SATORI and FIS. Interactions detected by FIS are sorted by frequency. Those detected by both methods are shown in blue, and ones detected only by FIS are shown in red. Top predictions are shown for the TAL-GATA dataset (a) the human promoter dataset (b), and the genomewide arabidopsis dataset (c). For each experiment, the 10 most frequent TF family interactions are shown in (d), (e), and (f) respectively.

Next, we compared the computation times for the two methods. As discussed earlier, unlike the FIS method, SATORI does not require re-calculation of the gradients to estimate the interactions, leading to much faster computation times: it processed all motif interactions 8 to 20 times faster than FIS (see Figure 6).

**Figure 6.**
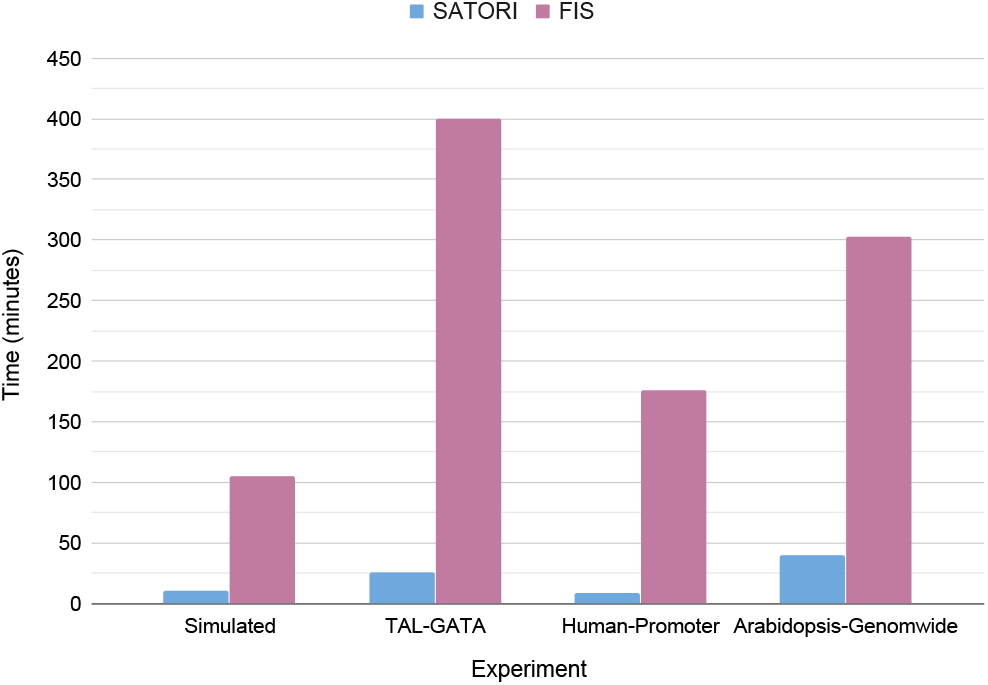
Run time in minutes for SATORI and FIS-based interaction estimation for the four datasets.

In summary, we find a very high level of overlap between the results of the two methods, which use very different approaches. This is important in view of the relatively small number of experimentally verified interactions that are available. Further wet-lab validation is needed to test the quality of the reported interactions by the two methods. High-frequency interactions consistently detected by both methods can be used as the most promising candidates for experimental follow-up.

## CONCLUSIONS AND FUTURE WORK

In this work we presented SATORI — a method for extracting interactions between the learned features of an attention-based deep learning model. Unlike existing methods, it only requires minimal post-processing and uses the sparsity of the attention matrix to infer the most salient interactions. We compared SATORI to the FIS interaction estimation method and reported a 10x speed-up in its computation time in most cases. Furthermore, the top predictions made by both methods show very high overlap, suggesting such interactions as promising targets for follow-up biological experiments. This high overlap, despite the big difference in the approach provides good evidence for their potential biological relevance.

The proposed method can be extended in several ways. In this work we focused on globally scoring interactions between TFs with known PWMs. This is in contrast to feature attribution methods that score the contribution of features in genomic regions of interest. We believe that the sparsity of the attention matrix could make it useful as an attribution method as well, but further experiments are required in order to validate that. SATORI is able to detect interactions between filters, even if they do not correspond to known TFs. Furthermore, the proposed methodology is flexible enough to be applied to deep networks that integrate multiple data modalities, and has potential applications outside of computational biology. For example, it can allow discovery of interactions between different characteristics of chromatin structure to provide a better understanding of the relationship between epigenetic markers such as histone modifications, DNA methylation, and nucleosome positioning and their contribution to the regulation of gene expression.

## AVAILABILITY

The source code for SATORI and the processed data and results are available at https://github.com/fahadahaf/satori.

