## Supplementary information for "A Self-Attention Model for Inferring Cooperativity between Regulatory Features"

### Supplementary Methods

#### Data collection and processing

##### A. Experiment 1: Simulated dataset

In this experiment, we simulated DNA sequences using random sampling from a distribution of [0.27, 0.23, 0.23, 0.27] for A, C, G, and T respectively as used for a similar dataset generated by Greenside et al. [1]. We generated 120,000 sequences each with a length of 200 bp. Similar to Greenside et al. [1], we randomly embedded instances of the motifs of both ELF1 and SIX5 transcription factors in 40,000 of the total sequences. This was our positive set of examples where we essentially simulated interactions between the aforementioned motifs. In the negative set (80,000 sequences), we embedded instances of either ELF1 or SIX5 in a sequence (but not both). Moreover, we embedded instances of the AP1 and TAL1 motifs across all examples. The motifs for the four transcription factors were obtained from Kheradpour et al. [2].

**TF Database information**

To map CNN filters to motifs of known TFs, we used TomTom with a custom TF database (MEME format) containing PWMs of the four transcription factors: SIX5, ELF1, AP1, and TAL1.

##### B. Experiment 2: TAL-GATA ChIP-peaks

Here we followed the same strategy described in DFIM [1]: ChIP-Seq peaks were downloaded for the three TFs TAL1, GATA1, and GATA2 from the ENCODE [4] database in the K562 cell line (hg19 genome assembly and annotations). For the chromatin accessibility data, we downloaded processed DNase I Hypersensitive Sites (DHSs) from the ENCODE database for the corresponding cell line. The raw data (bed files) is available at DFIM github repository. Next, every ChIP-Seq peak for the three transcription factors was searched for an overlap with DHSs in the K562 cell line. If an overlap was found, the sequence of the ChIP-Seq peak was extended 500 bp upstream and downstream from its center. This served as a positive set in our binary classification problem. For the negative set, we randomly sampled 80,000 examples from all K562 DHSs that didn't overlap a ChIP-Seq peak for any of the three transcription factors.

**TF Database information**

In this experiment, we used two TF databases: the first one was a custom motif file with PWMs of TAL1, GATA1, and GATA2 transcription factors. This was because we wanted to directly compare our model to DFIM [1] where Greenside et al. measured interactions between the aforementioned transcription factors. The second reference was the entire CISBP TF database [3] that we used in order to infer other TF interactions within the ChIP-Seq peaks.

##### C. Experiment 3: Human promoter DHSs

In this experiment, we used DHSs overlapping gene promoter regions across the entire human genome. We used the pipeline described by Kelley et al. in Basset [5]: DHSs were downloaded for 164 human immortalized cell lines from the ENCODE [4] and ROADMAP [6] consortia (more information on the samples can be found in their paper and accompanied supplement). These regions of open chromatin were merged if they overlapped more than 200 bp. Finally, every DHS was extended to a length of 600 bp around its center. Kelley et al. [5] used DHSs across the entire genome however, we selected only those which overlapped the human promoter regions. To do that, we defined promoter as a region of 1000 bp upstream of the transcription start site (TSS) of a gene—Ensemble based hg19/GrCh37 reference and annotations were used. The final dataset had 20,613 genomic sequences of the corresponding DHSs (that overlapped the human promoters). The targets in this case were either a single label or multiple labels, corresponding to the164 cell lines in which the DHSs were observed.

**TF Database information**
In motif analysis (and later in the TF interactions), we used the human CISBP transcription factor database [3].

##### D. Experiment 4: Genome-wide arabidopsis regions of open chromatin
Here we designed a similar experiment as described in the previous section. The dataset was constructed using the same procedure described above as used by Kelley et al. [5]. We used regions of open chromatin: DHSs and ATAC-Seq based Transposase Hypersensitive Sites (THSs), across the entire arabidopsis genome using TAIR10 annotations. We used the following publicly available datasets (GEO accession numbers provided):

1. **For DHSs:** GSE53322, GSE53324, GSE53323, GSE46987, GSE34318
2. **For THSs:** GSE89346, GSE85203, GSE101940, GSE116287, GSE101482

We ended up with 88,245 examples in our final dataset across 36 different samples. Note that peaks occurring in multiple biological samples were merged.

**TF Database information**
Here we used the DAP-Seq based arabidopsis transcription factors database [7].

#### Limitations of the TomTom motif comparison tool
We used the TomTom tool from the MEME suite [8] to map CNN filters to motifs of known transcription factors. In some cases, we observed the match to be dubious despite the tool assigning it a significant p-value. This is shown in Supplementary Figure F4 for two of our CNN filters matching the known TF motifs of HOXA2 and ZNF263 in the human CISBP database [3].
By default, TomTom uses *Pearson correlation* for comparing motifs. However, we obtained better results using the *Euclidean* distance.

#### Availability of TomTom results
For the four experiments, the TomTom motif matching results are available at our GitHub repository: https://github.com/fahadahaf/satori/tree/master/data/TomTom_files

### Details on the multi-head self-attention layer

The input to the multi-head self-attention (MHA) layer is a $d \times R$ matrix where $d$ is the length of the input sequence after the max-pooling operation—which is the last step in the convolutional layer—and $R$ is the dimensionality of the input which equals:

1. Number of CNN filters when the recurrent layer is absent (see hyperparameter *CNN_filters* in Table S1).
2. Size of the RNN hidden dimension when the recurrent layer is present (see hyperparameter *RNN_hiddensize* in Table S1).

For an individual head, the attention output (see *equation 8* in the main text) is a $d \times h$ matrix where $h$ is the size of the individual attention head, defined by the hyperparameter *singlehead_size* in Table S1. For $k$ individual attention heads (for the value of $k$, see hyperparameter *num_multiheads* in Table S1), we concatenate the outputs and end up with a $d \times kh$ matrix. Next, the concatenated matrix ($Z_m$) is linearly transformed using the following:

$$M = Z_m^{\top} W_m,$$

where $W_m$ is the associated weight matrix of dimensions $kh \times p$, and $p$ is the size of the multi-head linear transformation, defined by the hyperparameter *multihead_size* in Table S1.

Next, the matrix $M$ is summed across the columns and standardized, leading to a $p$-dimensional vector, which is passed to the fully connected layer. As mentioned in the main text, we find that this step is not only computationally efficient but also leads to better model accuracy in comparison to flattening the attention layer output. Note that the hyperparamter *readout_strategy* in Table S1 can be used to determine whether to do the aforementioned step (parameter value = "normalize") or flatten (parameter value = "flatten") the matrix $M$.

#### Supplementary Figures

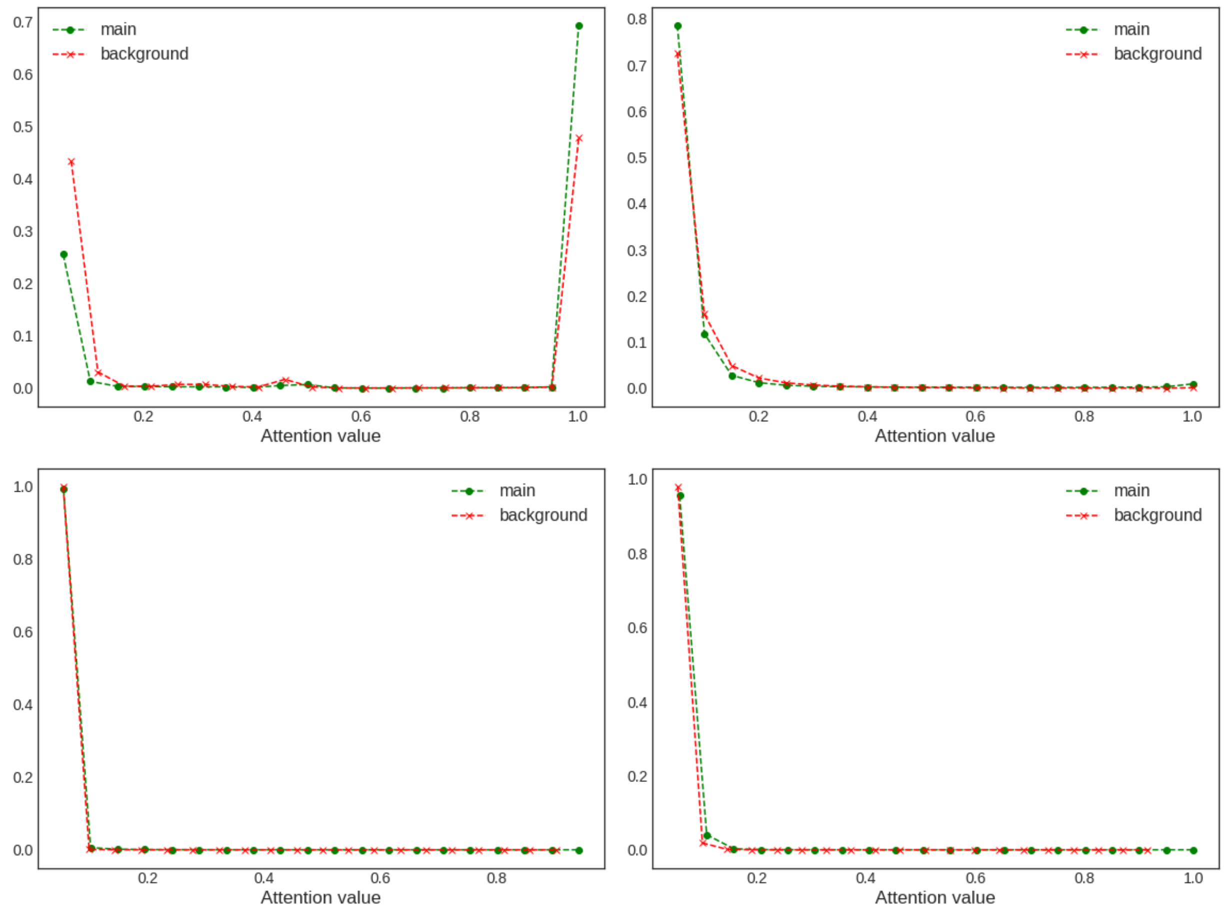


**Figure F1:** Distribution of the attention values for the test and the background sets are shown for the four experiments: simulated dataset (top-left), TAL-GATA ChIP peaks (top-right), human promoter DHSs (bottom-left) and arabidopsis accessible chromatin (bottom-right). The actual frequencies (y-axis) are normalized by total sizes of the test and background sets. This figure helps in selecting the appropriate attention cutoff, one of the parameters of SATORI. We use a default value of 0.10, although the model produces results at different thresholds.

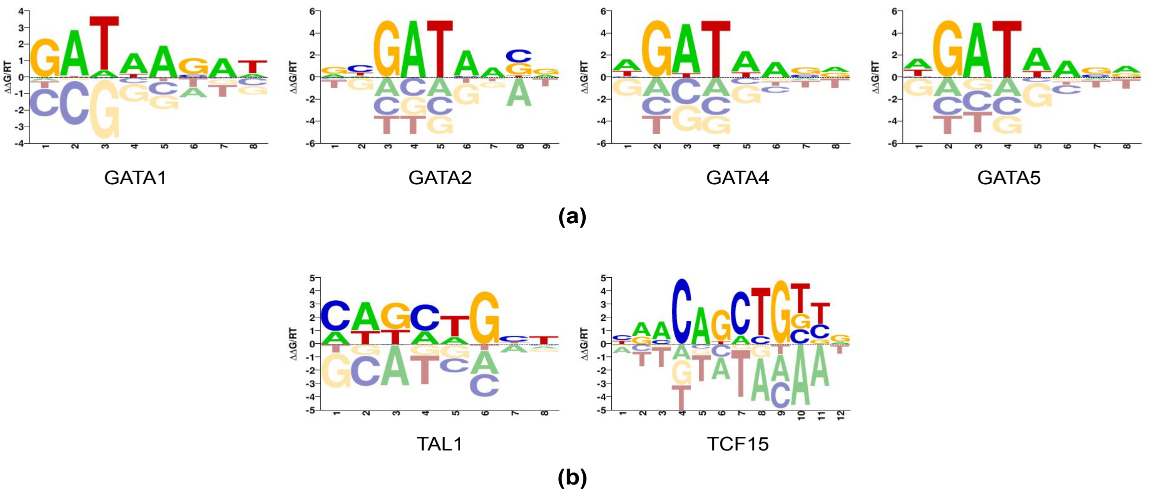


**Figure F2:** Similarities between motifs of GATA variants (a). Similarly, TAL1 and TCF15, both belonging to the bHLH family, have very similar motifs (CAGCTG consensus) (b).

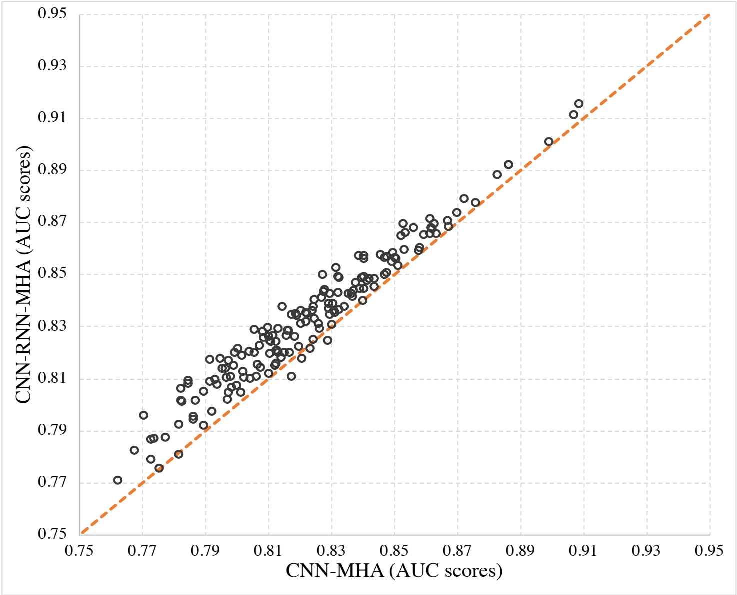


**Figure F3:** AUC scores for DHSs in human promoters across 164 cell types, achieved by the two model variants. Each circle represents performance on detecting DHSs in that cell line.

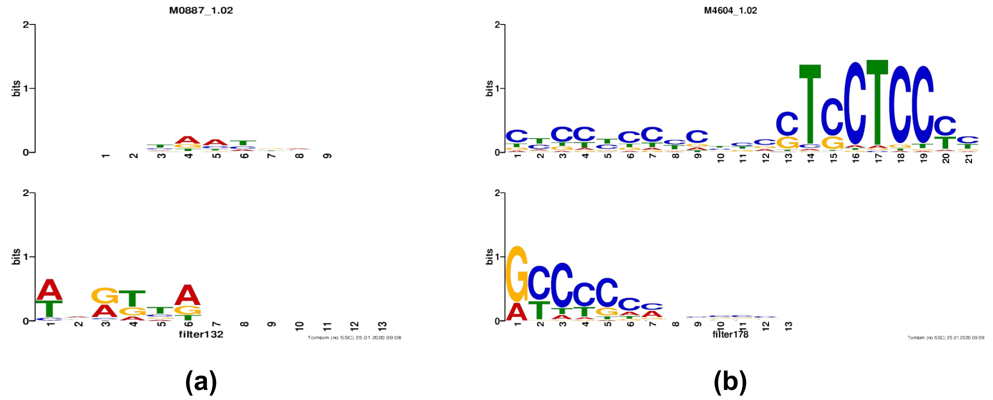


**Figure F4:** Limitations of the TomTom motif comparison tool. Matches shown here are detected as statistically significant (q-value < 0.01) for both (a) HOXA2 and (b) ZNF263. The top row depicts the gold standard motif in the CISBP database, and the bottom row shows the CNN filter/motif.

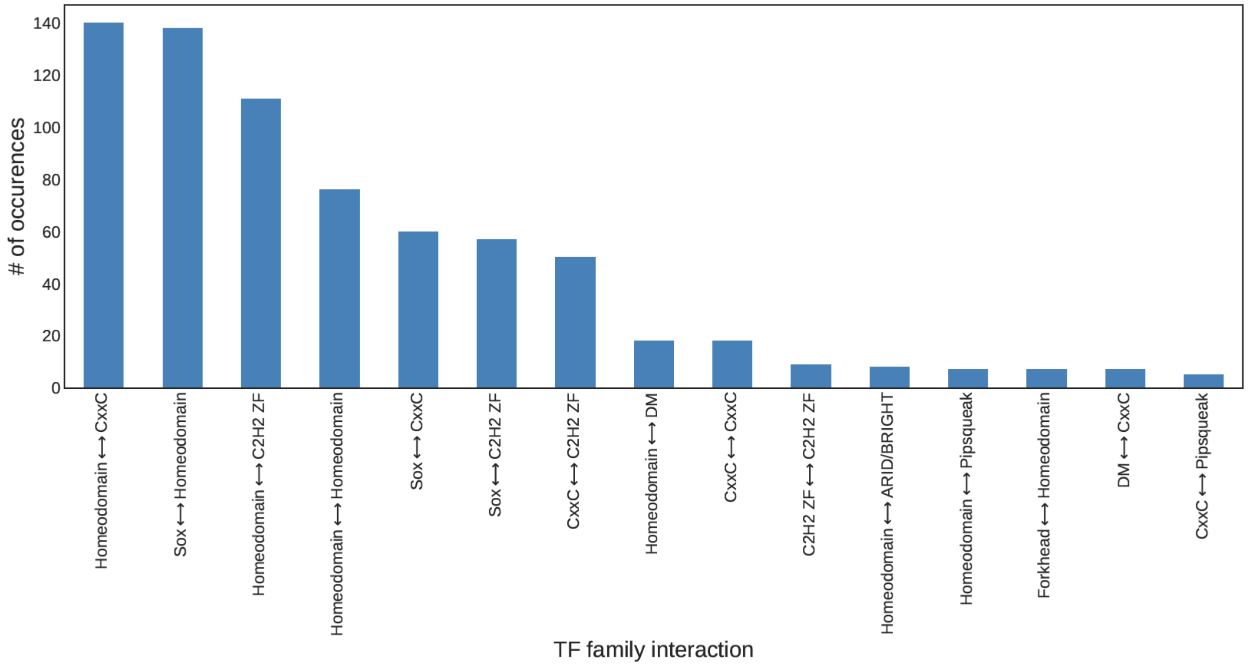


**Figure F5:** The most frequent interacting transcription factor families in human promoter regions.

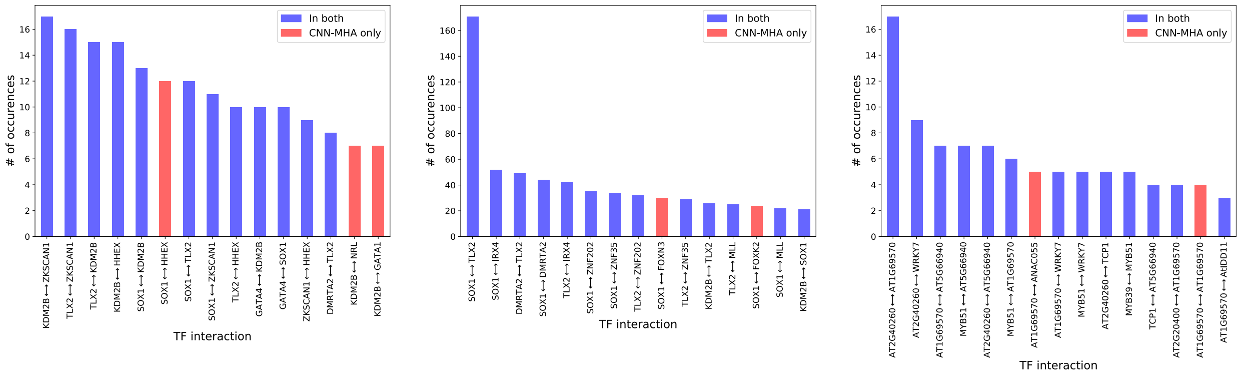


(a) (b) (c)

**Figure F6:** Common interactions in the top predictions of the two architectures: CNN-MHA and CNN-RNN-MHA. Interactions predicted using the CNN-MHA model are sorted by frequency. Those predicted by both methods are shown in blue, and ones predicted only by CNN-MHA are shown in red. Top predictions are shown for the TAL-GATA dataset (a) the human promoter dataset (b), and the genome-wide arabidopsis dataset (c).

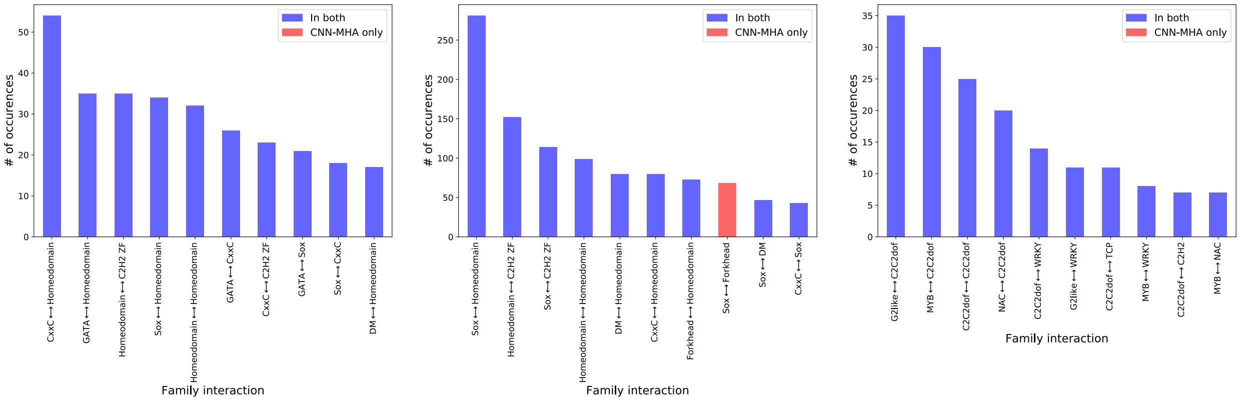


(a) (b) (c)

**Figure F7:** Common TF family interactions in the top predictions of the two architectures: CNN-MHA and CNN-RNN-MHA. Top predictions are shown for the TAL-GATA dataset (a) the human promoter dataset (b), and the genome-wide arabidopsis dataset (c).

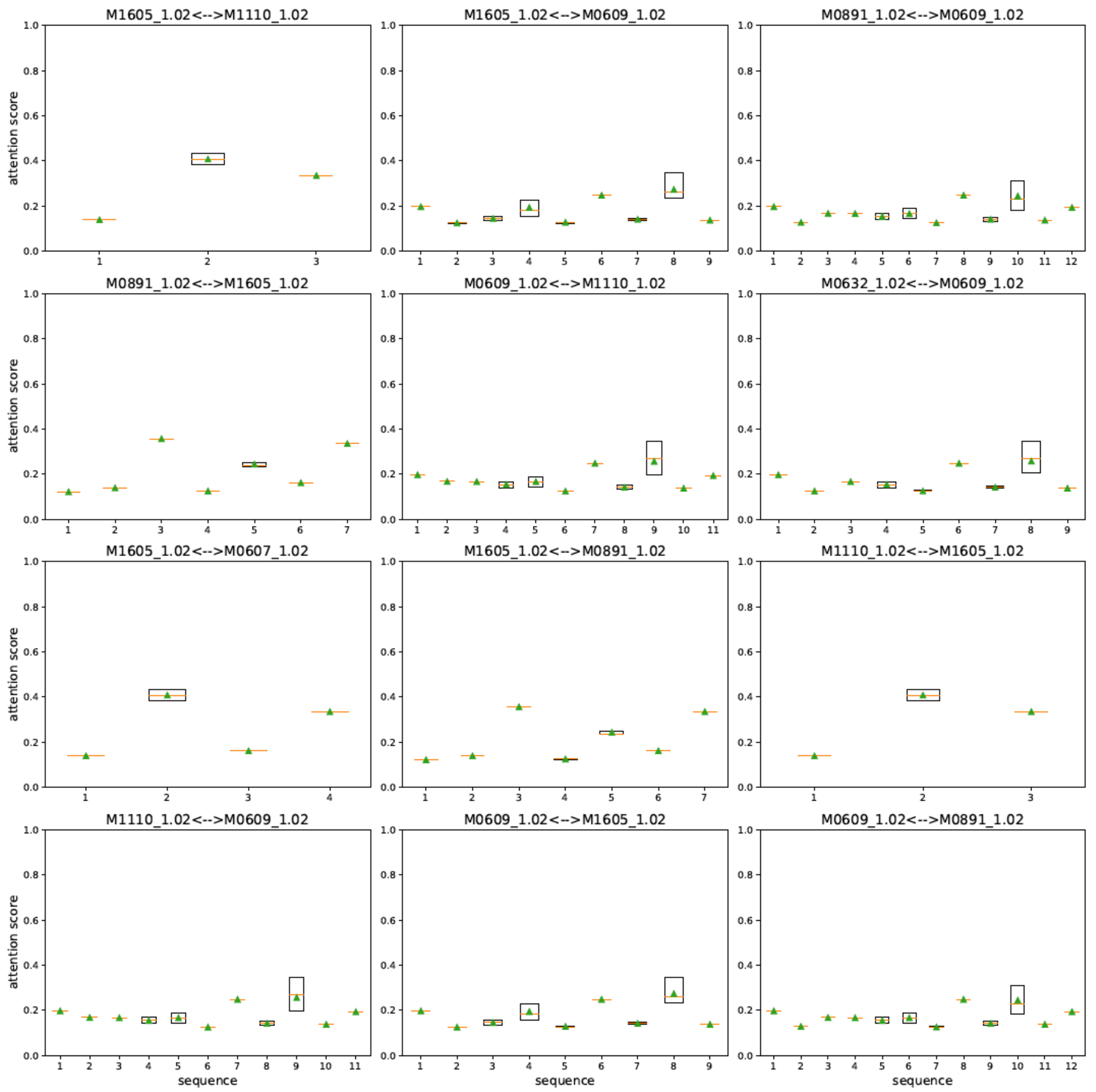


**Figure F8:** Distribution of attention scores (box plots) for twelve TF-TF interactions, each with multiple matching filters across all human promoter test sequences in which they appear.

### Supplementary Tables

**Table S1:** List of neural network hyperparameters common across the four experiments.

| Hyperparameter | type | Description |
|---|---|---|
| input_channels | int | Number of original input channels [Default: 4, for DNA] |
| num_multiheads | int | Number of heads in the multi-head attention [Default: 8] |
| singlehead_size | int | Size of the single attention head [Default: 32] |
| multihead_size | int | Size of the final attention after concatenation [Default: 100] |
| use_pooling | bool | Whether to use max pooling in the attention layer [Default: False] |
| pooling_val | int | Size of the max pooling window in the attention layer [Default: 6] |
| readout_strategy | str | Read-out strategy of the attention layer [Default: “normalize”] |
| use_RNN | bool | Whether to use the RNN layer [Default: True] |
| CNN1_useexponential | bool | Whether to use exponential activation in the convolution [Default: False] |
| CNN_filters | int | Number of filters in the convolutional layer [Default: 200] |
| CNN_filtersize | int | Filter size in the convolution [Default: 13] |
| use_CNNpool | bool | Whether to use max pooling in the convolutional layer [Default: True] |
| CNN_poolsize | int | Size of the max pooling window in the convolutional layer [Default: 6] |
| RNN_hiddensize | int | Size of the RNN/LSTM unit [Default: 100] |
| batch_size | int | Batch size in training and testing [Default: 256] |
| num_epochs | int | Number of training epochs [Default: 50] |

**Table S2:** Summary of all significant interactions in the simulated/toy dataset. Our model is able to recover multiple interactions involving SIX5 and ELF1 TF motifs. We also provide the actual CNN filter interactions in the first column, named based on the total number of filters in the convolutional layer.

| Filter interaction | TF motif interaction | Adjusted p-value |
|---|---|---|
| filter033⟷filter170 | SIX5⟷ELF1 | 1.47E-42 |
| filter170⟷filter181 | ELF1⟷SIX5 | 1.50E-39 |
| filter033⟷filter111 | SIX5⟷ELF1 | 1.12E-20 |
| filter111⟷filter181 | ELF1⟷SIX5 | 3.17E-22 |
| filter033⟷filter091 | SIX5⟷ELF1 | 2.44E-18 |
| filter091⟷filter181 | ELF1⟷SIX5 | 2.14E-12 |
| filter033⟷filter055 | SIX5⟷ELF1 | 2.64E-08 |
| filter055⟷filter181 | ELF1⟷SIX5 | 3.84E-04 |
| filter019⟷filter033 | ELF1⟷SIX5 | 1.35E-07 |
| filter019⟷filter181 | ELF1⟷SIX5 | 2.48E-12 |
| filter091⟷filter199 | ELF1⟷SIX5 | 2.82E-03 |

**Table S3:** All significant interactions between TAL1 and GATA transcription factors. Note that in this case, a custom TF database was used containing motifs for TAL1, GATA1, and GATA2. In case of TAL1, other TFs (LYL1, NHLH2, and TAL2) also shared the same binding site motif and hence are mentioned here.

| TF A | TF B | Adjusted p-value |
|---|---|---|
| GATA2 | LYL1, **TAL1**, NHLH2, TAL2 | 4.37E-24 |
| GATA2 | LYL1, **TAL1**, NHLH2, TAL2 | 8.18E-21 |
| LYL1, **TAL1**, NHLH2, TAL2 | GATA2 | 5.71E-19 |
| LYL1, **TAL1**, NHLH2, TAL2 | GATA2 | 1.54E-16 |
| LYL1, **TAL1**, NHLH2, TAL2 | GATA2 | 8.68E-11 |
| GATA2 | LYL1, **TAL1**, NHLH2, TAL2 | 3.92E-10 |
| LYL1, **TAL1**, NHLH2, TAL2 | GATA2 | 3.27E-08 |
| LYL1, **TAL1**, NHLH2, TAL2 | GATA2 | 3.28E-08 |
| GATA2 | LYL1, **TAL1**, NHLH2, TAL2 | 5.66E-05 |
| LYL1, **TAL1**, NHLH2, TAL2 | GATA2 | 6.76E-04 |
| LYL1, **TAL1**, NHLH2, TAL2 | GATA2 | 4.51E-03 |
| LYL1, **TAL1**, NHLH2, TAL2 | GATA2 | 4.66E-03 |
| LYL1, **TAL1**, NHLH2, TAL2 | GATA2 | 4.72E-03 |
| LYL1, **TAL1**, NHLH2, TAL2 | GATA2 | 5.41E-03 |

**Table S6:** A list of known TF interactions identified by our model in the human promoter dataset. TRRUSTv2 [9] and HIPPIE v2.0 [10] were used as references for known interactions. The level of significance (adjusted p-value) assigned by our model to each interaction is provided in the last column.

| Motif interaction | TF1 family | TF2 family | adjusted p-value | Database |
|---|---|---|---|---|
| DNMT1⟷LCOR | CxxC | Pipsqueak | 3.38E-29 | TRRUSTv2 |
| DNMT1⟷EGR1 | C2H2 ZF | CxxC | 1.30E-22 | |
| EGR1⟷LCOR | C2H2 ZF | Pipsqueak | 3.58E-08 | |
| FOXK2⟷EGR1 | C2H2 ZF | Forkhead | 1.09E-04 | |
| DNMT1⟷KDM2B | CxxC | CxxC | 1.61E-33 | HIPPIE v2.0 |
| DNMT1⟷LCOR | CxxC | Pipsqueak | 3.38E-29 | |
| HHEX⟷TLX2 | Homeodomain | Homeodomain | 5.88E-23 | |
| KDM2B⟷LCOR | CxxC | Pipsqueak | 2.36E-18 | |
| KDM2B⟷HHEX | Homeodomain | CxxC | 5.24E-18 | |
| FOXK2⟷DNMT1 | Forkhead | CxxC | 3.63E-12 | |
| HHEX⟷ZNF32 | Homeodomain | C2H2 ZF | 1.24E-07 | |
| SOX3⟷EGR1 | Sox | C2H2 ZF | 3.06E-07 | |
| FOXK2⟷KDM2B | Forkhead | CxxC | 2.07E-05 | |

**Table S7:** Summary of the number of unique statistically significant TF interactions reported by SATORI and FIS for the three real-world datasets.

| Experiment | SATORI | FIS |
|---|---|---|
| TAL-GATA ChIP-Seq | 172 | 284 |
| Human promoters | 123 | 168 |
| Arabidopsis genome-wide | 687 | 511 |

**Table S4:** A list of all unique TF interactions in the TAL-GATA ChIP-Seq peaks dataset. (appended to this supplement.)

**Table S5:** A list of all unique significant interactions in the human promoters. (appended to this supplement.)

| Table S4: TAL-GATA ChIP unique TF interactions | | | |
|---|---|---|---|
| filter_interaction | TF_Interaction | Family_Interaction | adjusted_pval |
| filter96⟷filter104 | DMRTA2⟷FOXN3 | DM⟷Forkhead | 0.016273919 |
| filter104⟷filter129 | DMRTA2⟷GATA5 | DM⟷GATA | 0.014186112 |
| filter30⟷filter133 | DMRTA2⟷GSX1 | Homeodomain⟷DM | 0.03069758 |
| filter62⟷filter104 | DMRTA2⟷HES2 | DM⟷bHLH | 0.002780668 |
| filter30⟷filter160 | DMRTA2⟷HHEX | Homeodomain⟷DM | 0.027453357 |
| filter104⟷filter153 | DMRTA2⟷HOXA11 | Homeodomain⟷DM | 0.004447103 |
| filter104⟷filter143 | DMRTA2⟷KDM2B | DM⟷CxxC | 5.79E-05 |
| filter104⟷filter199 | DMRTA2⟷PRKRIR | DM⟷THAP finger | 0.015684653 |
| filter30⟷filter57 | DMRTA2⟷RFX7 | DM⟷RFX | 0.038614307 |
| filter30⟷filter108 | DMRTA2⟷RXRB | DM⟷Nuclear receptor | 0.001270302 |
| filter30⟷filter159 | DMRTA2⟷SOX1 | DM⟷Sox | 0.032421057 |
| filter61⟷filter104 | DMRTA2⟷TCF15 | DM⟷bHLH | 0.005685621 |
| filter30⟷filter196 | DMRTA2⟷TLX2 | Homeodomain⟷DM | 0.026231892 |
| filter78⟷filter104 | DMRTA2⟷ZKSCAN1 | C2H2 ZF⟷DM | 0.04570174 |
| filter17⟷filter30 | DUXA⟷DMRTA2 | Homeodomain⟷DM | 0.002179452 |
| filter17⟷filter96 | DUXA⟷FOXN3 | Homeodomain⟷Forkhead | 7.49E-06 |
| filter17⟷filter129 | DUXA⟷GATA5 | Homeodomain⟷GATA | 0.008553442 |
| filter17⟷filter133 | DUXA⟷GSX1 | Homeodomain⟷Homeodomain | 0.001298725 |
| filter17⟷filter62 | DUXA⟷HES2 | Homeodomain⟷bHLH | 1.66E-05 |
| filter17⟷filter160 | DUXA⟷HHEX | Homeodomain⟷Homeodomain | 0.0025193 |
| filter17⟷filter153 | DUXA⟷HOXA11 | Homeodomain⟷Homeodomain | 0.008288421 |
| filter17⟷filter27 | DUXA⟷IRX4 | Homeodomain⟷Homeodomain | 2.63E-05 |
| filter17⟷filter143 | DUXA⟷KDM2B | Homeodomain⟷CxxC | 2.65E-10 |
| filter17⟷filter199 | DUXA⟷PRKRIR | Homeodomain⟷THAP finger | 0.002570267 |
| filter17⟷filter108 | DUXA⟷RXRB | Homeodomain⟷Nuclear receptor | 0.030577167 |
| filter17⟷filter171 | DUXA⟷SOX1 | Homeodomain⟷Sox | 0.012312924 |
| filter17⟷filter61 | DUXA⟷TCF15 | Homeodomain⟷bHLH | 1.93E-05 |
| filter17⟷filter152 | DUXA⟷TLX2 | Homeodomain⟷Homeodomain | 0.024489316 |
| filter17⟷filter78 | DUXA⟷ZKSCAN1 | Homeodomain⟷C2H2 ZF | 0.001731158 |
| filter17⟷filter86 | DUXA⟷ZNF202 | Homeodomain⟷C2H2 ZF | 0.014991929 |
| filter96⟷filter133 | FOXN3⟷GSX1 | Homeodomain⟷Forkhead | 0.004969243 |
| filter96⟷filter153 | FOXN3⟷HOXA11 | Homeodomain⟷Forkhead | 1.32E-05 |
| filter96⟷filter143 | FOXN3⟷KDM2B | Forkhead⟷CxxC | 0.000622207 |
| filter96⟷filter199 | FOXN3⟷PRKRIR | Forkhead⟷THAP finger | 0.001806896 |
| filter96⟷filter108 | FOXN3⟷RXRB | Forkhead⟷Nuclear receptor | 0.019092169 |
| filter96⟷filter159 | FOXN3⟷SOX1 | Forkhead⟷Sox | 0.00264225 |
| filter96⟷filter129 | GATA5⟷FOXN3 | GATA⟷Forkhead | 3.41E-07 |
| filter129⟷filter133 | GATA5⟷GSX1 | Homeodomain⟷GATA | 0.004665757 |
| filter129⟷filter160 | GATA5⟷HHEX | Homeodomain⟷GATA | 0.011465718 |
| filter129⟷filter153 | GATA5⟷HOXA11 | Homeodomain⟷GATA | 3.06E-05 |
| filter129⟷filter198 | GATA5⟷PAX2 | GATA⟷Homeodomain,Paired box | 0.010233609 |
| filter89⟷filter129 | GATA5⟷RARB | GATA⟷Nuclear receptor | 0.037681423 |
| filter84⟷filter139 | GATA5⟷RARG | GATA⟷Nuclear receptor | 1.03E-12 |

| | | | |
|---|---|---|---|
| filter108⟷filter129 | GATA5⟷RXRB | GATA⟷Nuclear receptor | 0.007053379 |
| filter84⟷filter154 | GATA5⟷UNKNOWN | GATA⟷UNKNOWN | 0.01304683 |
| filter133⟷filter153 | GSX1⟷HOXA11 | Homeodomain⟷Homeodomain | 0.002127705 |
| filter133⟷filter143 | GSX1⟷KDM2B | Homeodomain⟷CxxC | 5.75E-06 |
| filter133⟷filter199 | GSX1⟷PRKRIR | Homeodomain⟷THAP finger | 8.59E-06 |
| filter133⟷filter171 | GSX1⟷SOX1 | Homeodomain⟷Sox | 0.000746888 |
| filter62⟷filter96 | HES2⟷FOXN3 | bHLH⟷Forkhead | 1.48E-05 |
| filter62⟷filter129 | HES2⟷GATA5 | bHLH⟷GATA | 2.18E-10 |
| filter62⟷filter133 | HES2⟷GSX1 | Homeodomain⟷bHLH | 0.002993578 |
| filter62⟷filter160 | HES2⟷HHEX | Homeodomain⟷bHLH | 0.025163548 |
| filter62⟷filter153 | HES2⟷HOXA11 | Homeodomain⟷bHLH | 1.35E-08 |
| filter62⟷filter175 | HES2⟷KDM2B | bHLH⟷CxxC | 0.049123556 |
| filter62⟷filter199 | HES2⟷PRKRIR | bHLH⟷THAP finger | 8.52E-08 |
| filter62⟷filter108 | HES2⟷RXRB | bHLH⟷Nuclear receptor | 0.006442359 |
| filter62⟷filter159 | HES2⟷SOX1 | bHLH⟷Sox | 0.003489763 |
| filter62⟷filter78 | HES2⟷ZKSCAN1 | C2H2 ZF⟷bHLH | 4.53E-08 |
| filter160⟷filter199 | HHEX⟷PRKRIR | Homeodomain⟷THAP finger | 0.025461016 |
| filter160⟷filter167 | HHEX⟷SOX1 | Homeodomain⟷Sox | 1.29E-05 |
| filter153⟷filter160 | HOXA11⟷HHEX | Homeodomain⟷Homeodomain | 0.000478357 |
| filter153⟷filter199 | HOXA11⟷PRKRIR | Homeodomain⟷THAP finger | 6.69E-09 |
| filter153⟷filter159 | HOXA11⟷SOX1 | Homeodomain⟷Sox | 9.52E-05 |
| filter27⟷filter30 | IRX4⟷DMRTA2 | Homeodomain⟷DM | 4.27E-05 |
| filter27⟷filter96 | IRX4⟷FOXN3 | Homeodomain⟷Forkhead | 4.17E-08 |
| filter27⟷filter129 | IRX4⟷GATA5 | Homeodomain⟷GATA | 0.001458864 |
| filter27⟷filter133 | IRX4⟷GSX1 | Homeodomain⟷Homeodomain | 0.00010058 |
| filter1⟷filter62 | IRX4⟷HES2 | Homeodomain⟷bHLH | 0.028695778 |
| filter27⟷filter160 | IRX4⟷HHEX | Homeodomain⟷Homeodomain | 0.006852958 |
| filter1⟷filter153 | IRX4⟷HOXA11 | Homeodomain⟷Homeodomain | 0.004003367 |
| filter1⟷filter143 | IRX4⟷KDM2B | Homeodomain⟷CxxC | 0.02803546 |
| filter27⟷filter198 | IRX4⟷PAX2 | Homeodomain⟷Homeodomain,Paired box | 0.000599869 |
| filter1⟷filter199 | IRX4⟷PRKRIR | Homeodomain⟷THAP finger | 0.015852369 |
| filter27⟷filter89 | IRX4⟷RARB | Homeodomain⟷Nuclear receptor | 0.000154095 |
| filter27⟷filter139 | IRX4⟷RARG | Homeodomain⟷Nuclear receptor | 1.70E-05 |
| filter27⟷filter57 | IRX4⟷RFX7 | Homeodomain⟷RFX | 1.55E-05 |
| filter27⟷filter108 | IRX4⟷RXRB | Homeodomain⟷Nuclear receptor | 0.000122858 |
| filter27⟷filter35 | IRX4⟷SIX4 | Homeodomain⟷Homeodomain | 0.036570768 |
| filter1⟷filter171 | IRX4⟷SOX1 | Homeodomain⟷Sox | 0.044628918 |
| filter1⟷filter61 | IRX4⟷TCF15 | Homeodomain⟷bHLH | 0.01621677 |
| filter27⟷filter189 | IRX4⟷ZKSCAN1 | Homeodomain⟷C2H2 ZF | 0.044039359 |
| filter47⟷filter84 | KDM2B⟷GATA5 | CxxC⟷GATA | 0.013086664 |
| filter143⟷filter160 | KDM2B⟷HHEX | Homeodomain⟷CxxC | 0.001874772 |
| filter143⟷filter153 | KDM2B⟷HOXA11 | Homeodomain⟷CxxC | 3.19E-09 |
| filter175⟷filter199 | KDM2B⟷PRKRIR | CxxC⟷THAP finger | 0.029126468 |
| filter143⟷filter159 | KDM2B⟷SOX1 | CxxC⟷Sox | 0.00015308 |
| filter67⟷filter84 | PRKRIR⟷GATA5 | THAP finger⟷GATA | 7.18E-05 |

| | | | |
|---|---|---|---|
| filter57⟷filter96 | RFX7⟷FOXN3 | RFX⟷Forkhead | 0.003548686 |
| filter57⟷filter129 | RFX7⟷GATA5 | RFX⟷GATA | 0.002120495 |
| filter57⟷filter62 | RFX7⟷HES2 | RFX⟷bHLH | 1.43E-05 |
| filter57⟷filter153 | RFX7⟷HOXA11 | Homeodomain⟷RFX | 0.00190261 |
| filter57⟷filter175 | RFX7⟷KDM2B | RFX⟷CxxC | 0.020444823 |
| filter57⟷filter199 | RFX7⟷PRKRIR | RFX⟷THAP finger | 4.14E-06 |
| filter57⟷filter108 | RFX7⟷RXRB | RFX⟷Nuclear receptor | 0.00589543 |
| filter57⟷filter159 | RFX7⟷SOX1 | RFX⟷Sox | 0.023260037 |
| filter57⟷filter61 | RFX7⟷TCF15 | RFX⟷bHLH | 0.01193643 |
| filter57⟷filter189 | RFX7⟷ZKSCAN1 | RFX⟷C2H2 ZF | 0.01226628 |
| filter57⟷filter86 | RFX7⟷ZNF202 | RFX⟷C2H2 ZF | 0.014782827 |
| filter108⟷filter153 | RXRB⟷HOXA11 | Homeodomain⟷Nuclear receptor | 3.88E-06 |
| filter108⟷filter143 | RXRB⟷KDM2B | Nuclear receptor⟷CxxC | 6.81E-05 |
| filter108⟷filter199 | RXRB⟷PRKRIR | Nuclear receptor⟷THAP finger | 0.000642153 |
| filter108⟷filter167 | RXRB⟷SOX1 | Nuclear receptor⟷Sox | 1.22E-05 |
| filter10⟷filter104 | SIX4⟷DMRTA2 | Homeodomain⟷DM | 0.004236246 |
| filter10⟷filter80 | SIX4⟷DNMT1 | Homeodomain⟷CxxC | 0.029940625 |
| filter10⟷filter17 | SIX4⟷DUXA | Homeodomain⟷Homeodomain | 7.44E-05 |
| filter10⟷filter96 | SIX4⟷FOXN3 | Homeodomain⟷Forkhead | 1.06E-10 |
| filter35⟷filter84 | SIX4⟷GATA5 | Homeodomain⟷GATA | 0.009772488 |
| filter10⟷filter133 | SIX4⟷GSX1 | Homeodomain⟷Homeodomain | 1.17E-08 |
| filter10⟷filter62 | SIX4⟷HES2 | Homeodomain⟷bHLH | 9.62E-15 |
| filter10⟷filter160 | SIX4⟷HHEX | Homeodomain⟷Homeodomain | 5.49E-12 |
| filter10⟷filter153 | SIX4⟷HOXA11 | Homeodomain⟷Homeodomain | 6.55E-17 |
| filter10⟷filter47 | SIX4⟷KDM2B | Homeodomain⟷CxxC | 0.039227817 |
| filter10⟷filter198 | SIX4⟷PAX2 | Homeodomain⟷Homeodomain,Paired box | 8.25E-06 |
| filter10⟷filter67 | SIX4⟷PRKRIR | Homeodomain⟷THAP finger | 0.000821242 |
| filter10⟷filter89 | SIX4⟷RARB | Homeodomain⟷Nuclear receptor | 0.000197954 |
| filter10⟷filter139 | SIX4⟷RARG | Homeodomain⟷Nuclear receptor | 0.000161492 |
| filter10⟷filter57 | SIX4⟷RFX7 | Homeodomain⟷RFX | 3.80E-06 |
| filter10⟷filter108 | SIX4⟷RXRB | Homeodomain⟷Nuclear receptor | 2.94E-10 |
| filter10⟷filter23 | SIX4⟷SOX1 | Homeodomain⟷Sox | 0.025635601 |
| filter10⟷filter63 | SIX4⟷TCF15 | Homeodomain⟷bHLH | 0.000833746 |
| filter10⟷filter152 | SIX4⟷TLX2 | Homeodomain⟷Homeodomain | 0.007335701 |
| filter10⟷filter154 | SIX4⟷UNKNOWN | Homeodomain⟷UNKNOWN | 0.000148137 |
| filter10⟷filter54 | SIX4⟷ZFX | Homeodomain⟷C2H2 ZF | 0.004371377 |
| filter10⟷filter41 | SIX4⟷ZKSCAN1 | Homeodomain⟷C2H2 ZF | 0.004145332 |
| filter5⟷filter84 | SNAI3⟷GATA5 | C2H2 ZF⟷GATA | 6.57E-06 |
| filter5⟷filter27 | SNAI3⟷IRX4 | Homeodomain⟷C2H2 ZF | 0.026424291 |
| filter5⟷filter10 | SNAI3⟷SIX4 | Homeodomain⟷C2H2 ZF | 0.001231352 |
| filter23⟷filter84 | SOX1⟷GATA5 | Sox⟷GATA | 0.014882266 |
| filter159⟷filter199 | SOX1⟷PRKRIR | Sox⟷THAP finger | 0.005266152 |
| filter61⟷filter96 | TCF15⟷FOXN3 | bHLH⟷Forkhead | 1.94E-07 |
| filter61⟷filter129 | TCF15⟷GATA5 | bHLH⟷GATA | 7.50E-06 |
| filter61⟷filter133 | TCF15⟷GSX1 | Homeodomain⟷bHLH | 3.17E-05 |

| | | | |
|---|---|---|---|
| filter62⟷filter63 | TCF15⟷HES2 | bHLH⟷bHLH | 0.015057894 |
| filter61⟷filter160 | TCF15⟷HHEX | Homeodomain⟷bHLH | 0.007053379 |
| filter63⟷filter153 | TCF15⟷HOXA11 | Homeodomain⟷bHLH | 0.015061257 |
| filter61⟷filter143 | TCF15⟷KDM2B | bHLH⟷CxxC | 8.97E-10 |
| filter61⟷filter199 | TCF15⟷PRKRIR | bHLH⟷THAP finger | 2.49E-06 |
| filter61⟷filter108 | TCF15⟷RXRB | bHLH⟷Nuclear receptor | 0.003760887 |
| filter61⟷filter159 | TCF15⟷SOX1 | bHLH⟷Sox | 0.024715282 |
| filter129⟷filter152 | TLX2⟷GATA5 | Homeodomain⟷GATA | 0.049552316 |
| filter16⟷filter62 | TLX2⟷HES2 | Homeodomain⟷bHLH | 0.005464262 |
| filter153⟷filter196 | TLX2⟷HOXA11 | Homeodomain⟷Homeodomain | 0.022440048 |
| filter27⟷filter173 | TLX2⟷IRX4 | Homeodomain⟷Homeodomain | 0.048355552 |
| filter16⟷filter143 | TLX2⟷KDM2B | Homeodomain⟷CxxC | 0.003728609 |
| filter173⟷filter199 | TLX2⟷PRKRIR | Homeodomain⟷THAP finger | 0.032834049 |
| filter152⟷filter167 | TLX2⟷SOX1 | Homeodomain⟷Sox | 0.017990916 |
| filter16⟷filter61 | TLX2⟷TCF15 | Homeodomain⟷bHLH | 0.003050355 |
| filter78⟷filter100 | TLX2⟷ZKSCAN1 | Homeodomain⟷C2H2 ZF | 0.005676285 |
| filter54⟷filter84 | ZFX⟷GATA5 | C2H2 ZF⟷GATA | 0.000126567 |
| filter78⟷filter96 | ZKSCAN1⟷FOXN3 | C2H2 ZF⟷Forkhead | 1.84E-05 |
| filter129⟷filter189 | ZKSCAN1⟷GATA5 | C2H2 ZF⟷GATA | 0.020347564 |
| filter78⟷filter133 | ZKSCAN1⟷GSX1 | Homeodomain⟷C2H2 ZF | 0.004532009 |
| filter78⟷filter160 | ZKSCAN1⟷HHEX | Homeodomain⟷C2H2 ZF | 0.000224975 |
| filter41⟷filter153 | ZKSCAN1⟷HOXA11 | Homeodomain⟷C2H2 ZF | 0.002052756 |
| filter41⟷filter143 | ZKSCAN1⟷KDM2B | C2H2 ZF⟷CxxC | 0.0373158 |
| filter189⟷filter199 | ZKSCAN1⟷PRKRIR | C2H2 ZF⟷THAP finger | 0.000486019 |
| filter78⟷filter108 | ZKSCAN1⟷RXRB | C2H2 ZF⟷Nuclear receptor | 0.000848699 |
| filter78⟷filter166 | ZKSCAN1⟷SOX1 | C2H2 ZF⟷Sox | 0.02176073 |
| filter63⟷filter78 | ZKSCAN1⟷TCF15 | C2H2 ZF⟷bHLH | 0.031414495 |
| filter78⟷filter90 | ZKSCAN1⟷ZNF202 | C2H2 ZF⟷C2H2 ZF | 0.009838888 |
| filter2⟷filter30 | ZNF202⟷DMRTA2 | C2H2 ZF⟷DM | 0.035636216 |
| filter86⟷filter96 | ZNF202⟷FOXN3 | C2H2 ZF⟷Forkhead | 0.02631752 |
| filter33⟷filter129 | ZNF202⟷GATA5 | C2H2 ZF⟷GATA | 0.018945943 |
| filter2⟷filter62 | ZNF202⟷HES2 | C2H2 ZF⟷bHLH | 0.007104995 |
| filter90⟷filter153 | ZNF202⟷HOXA11 | Homeodomain⟷C2H2 ZF | 0.010360849 |
| filter27⟷filter144 | ZNF202⟷IRX4 | Homeodomain⟷C2H2 ZF | 0.044577055 |
| filter90⟷filter143 | ZNF202⟷KDM2B | C2H2 ZF⟷CxxC | 0.01087048 |
| filter90⟷filter199 | ZNF202⟷PRKRIR | C2H2 ZF⟷THAP finger | 0.005676285 |
| filter10⟷filter148 | ZNF202⟷SIX4 | Homeodomain⟷C2H2 ZF | 0.049931244 |
| filter86⟷filter159 | ZNF202⟷SOX1 | C2H2 ZF⟷Sox | 0.046686771 |
| filter2⟷filter61 | ZNF202⟷TCF15 | C2H2 ZF⟷bHLH | 0.011246917 |

| Table S5: Human promoter unique TF interactions | | | |
|---|---|---|---|
| **filter_interaction** | **TF_Interaction** | **Family_Interaction** | **adjusted_pval** |
| filter133⟷filter178 | ARID3B⟷DNMT1 | ARID/BRIGHT⟷CxxC | 6.86E-12 |
| filter124⟷filter133 | DMRTA2⟷DNMT1 | DM⟷CxxC | 1.11E-14 |
| filter39⟷filter161 | DMRTA2⟷EGR1 | DM⟷C2H2 ZF | 9.98E-06 |
| filter101⟷filter143 | DMRTA2⟷HHEX | Homeodomain⟷DM | 5.71E-05 |
| filter161⟷filter162 | DNMT1⟷EGR1 | CxxC⟷C2H2 ZF | 0.000187249 |
| filter28⟷filter174 | DNMT1⟷HOXA11 | Homeodomain⟷CxxC | 4.29E-07 |
| filter155⟷filter159 | DNMT1⟷KDM2B | CxxC⟷CxxC | 0.005490864 |
| filter151⟷filter162 | DNMT1⟷LCOR | CxxC⟷Pipsqueak | 0.000117664 |
| filter133⟷filter148 | DNMT1⟷TET1 | CxxC⟷CxxC | 4.65E-33 |
| filter28⟷filter57 | DNMT1⟷ZNF202 | CxxC⟷C2H2 ZF | 0.000916891 |
| filter54⟷filter195 | DNMT1⟷ZNF263 | CxxC⟷C2H2 ZF | 0.002379617 |
| filter133⟷filter192 | DNMT1⟷ZNF35 | CxxC⟷C2H2 ZF | 9.40E-13 |
| filter71⟷filter107 | EGR1⟷ARID3B | C2H2 ZF⟷ARID/BRIGHT | 3.01E-05 |
| filter71⟷filter92 | EGR1⟷HMG20B | Sox⟷C2H2 ZF | 1.24E-06 |
| filter161⟷filter174 | EGR1⟷HOXA11 | Homeodomain⟷C2H2 ZF | 4.57E-06 |
| filter90⟷filter161 | EGR1⟷KDM2B | CxxC⟷C2H2 ZF | 0.001643881 |
| filter151⟷filter161 | EGR1⟷LCOR | C2H2 ZF⟷Pipsqueak | 1.31E-06 |
| filter131⟷filter161 | EGR1⟷SNAI3 | C2H2 ZF⟷C2H2 ZF | 5.24E-08 |
| filter71⟷filter148 | EGR1⟷TET1 | CxxC⟷C2H2 ZF | 2.15E-11 |
| filter71⟷filter195 | EGR1⟷ZNF263 | C2H2 ZF⟷C2H2 ZF | 0.002039419 |
| filter71⟷filter118 | EGR1⟷ZNF32 | C2H2 ZF⟷C2H2 ZF | 3.34E-08 |
| filter78⟷filter101 | ENSG00000235187⟷HHEX | Homeodomain⟷Ets | 6.55E-06 |
| filter17⟷filter133 | FOXK2⟷DNMT1 | Forkhead⟷CxxC | 5.90E-11 |
| filter17⟷filter161 | FOXK2⟷EGR1 | Forkhead⟷C2H2 ZF | 0.019849637 |
| filter17⟷filter101 | FOXK2⟷HHEX | Forkhead⟷Homeodomain | 8.81E-06 |
| filter80⟷filter138 | FOXK2⟷KDM2B | Forkhead⟷CxxC | 9.54E-05 |
| filter21⟷filter80 | FOXK2⟷TLX2 | Forkhead⟷Homeodomain | 0.01682431 |
| filter101⟷filter107 | HHEX⟷ARID3B | Homeodomain⟷ARID/BRIGHT | 0.000772337 |
| filter162⟷filter197 | HHEX⟷DNMT1 | Homeodomain⟷CxxC | 0.005126141 |
| filter10⟷filter71 | HHEX⟷EGR1 | Homeodomain⟷C2H2 ZF | 0.000478663 |
| filter101⟷filter174 | HHEX⟷HOXA11 | Homeodomain⟷Homeodomain | 1.44E-10 |
| filter151⟷filter197 | HHEX⟷LCOR | Homeodomain⟷Pipsqueak | 2.19E-10 |
| filter131⟷filter135 | HHEX⟷SNAI3 | Homeodomain⟷C2H2 ZF | 6.81E-09 |
| filter101⟷filter148 | HHEX⟷TET1 | Homeodomain⟷CxxC | 5.55E-24 |
| filter10⟷filter21 | HHEX⟷TLX2 | Homeodomain⟷Homeodomain | 0.022836552 |
| filter10⟷filter102 | HHEX⟷ZNF263 | Homeodomain⟷C2H2 ZF | 3.84E-05 |
| filter101⟷filter118 | HHEX⟷ZNF32 | Homeodomain⟷C2H2 ZF | 1.24E-07 |
| filter135⟷filter192 | HHEX⟷ZNF35 | Homeodomain⟷C2H2 ZF | 1.34E-06 |
| filter92⟷filter133 | HMG20B⟷DNMT1 | Sox⟷CxxC | 7.89E-20 |
| filter92⟷filter101 | HMG20B⟷HHEX | Sox⟷Homeodomain | 2.20E-07 |
| filter92⟷filter102 | HMG20B⟷ZNF263 | Sox⟷C2H2 ZF | 1.43E-06 |
| filter1⟷filter39 | HOXA6⟷DMRTA2 | Homeodomain⟷DM | 0.000279832 |
| filter1⟷filter54 | HOXA6⟷DNMT1 | Homeodomain⟷CxxC | 0.004288393 |

| | | | |
|---|---|---|---|
| filter1⟷filter161 | HOXA6⟷EGR1 | Homeodomain⟷C2H2 ZF | 1.27E-05 |
| filter1⟷filter78 | HOXA6⟷ENSG00000235187 | Homeodomain⟷Ets | 0.00691044 |
| filter1⟷filter197 | HOXA6⟷HHEX | Homeodomain⟷Homeodomain | 5.43E-06 |
| filter1⟷filter98 | HOXA6⟷KDM2B | Homeodomain⟷CxxC | 0.007277367 |
| filter1⟷filter151 | HOXA6⟷LCOR | Homeodomain⟷Pipsqueak | 1.50E-07 |
| filter1⟷filter128 | HOXA6⟷LHX5 | Homeodomain⟷Homeodomain | 0.01607373 |
| filter1⟷filter131 | HOXA6⟷SNAI3 | Homeodomain⟷C2H2 ZF | 0.002589448 |
| filter1⟷filter189 | HOXA6⟷SOX1 | Sox⟷Homeodomain | 0.011781033 |
| filter1⟷filter126 | HOXA6⟷TLX2 | Homeodomain⟷Homeodomain | 0.029590696 |
| filter1⟷filter59 | HOXA6⟷ZKSCAN1 | Homeodomain⟷C2H2 ZF | 0.002400118 |
| filter1⟷filter57 | HOXA6⟷ZNF202 | Homeodomain⟷C2H2 ZF | 1.16E-05 |
| filter1⟷filter195 | HOXA6⟷ZNF263 | Homeodomain⟷C2H2 ZF | 0.000182924 |
| filter1⟷filter118 | HOXA6⟷ZNF32 | Homeodomain⟷C2H2 ZF | 0.0009448 |
| filter33⟷filter133 | IRF3⟷DNMT1 | IRF⟷CxxC | 2.70E-13 |
| filter33⟷filter71 | IRF3⟷EGR1 | IRF⟷C2H2 ZF | 4.36E-07 |
| filter33⟷filter101 | IRF3⟷HHEX | IRF⟷Homeodomain | 7.17E-06 |
| filter33⟷filter130 | IRF3⟷TLX2 | IRF⟷Homeodomain | 4.30E-07 |
| filter138⟷filter178 | KDM2B⟷ARID3B | ARID/BRIGHT⟷CxxC | 0.000370208 |
| filter124⟷filter138 | KDM2B⟷DMRTA2 | DM⟷CxxC | 2.27E-05 |
| filter173⟷filter197 | KDM2B⟷HHEX | Homeodomain⟷CxxC | 0.00165582 |
| filter151⟷filter159 | KDM2B⟷LCOR | CxxC⟷Pipsqueak | 1.33E-07 |
| filter173⟷filter195 | KDM2B⟷ZNF263 | CxxC⟷C2H2 ZF | 0.038854905 |
| filter138⟷filter192 | KDM2B⟷ZNF35 | CxxC⟷C2H2 ZF | 1.34E-05 |
| filter121⟷filter133 | LHX5⟷DNMT1 | Homeodomain⟷CxxC | 3.80E-12 |
| filter71⟷filter121 | LHX5⟷EGR1 | Homeodomain⟷C2H2 ZF | 8.58E-06 |
| filter19⟷filter101 | LHX5⟷HHEX | Homeodomain⟷Homeodomain | 0.004020722 |
| filter21⟷filter83 | LHX5⟷TLX2 | Homeodomain⟷Homeodomain | 0.028737386 |
| filter41⟷filter102 | LHX5⟷ZNF263 | Homeodomain⟷C2H2 ZF | 5.73E-06 |
| filter131⟷filter133 | SNAI3⟷DNMT1 | CxxC⟷C2H2 ZF | 5.36E-30 |
| filter131⟷filter138 | SNAI3⟷KDM2B | CxxC⟷C2H2 ZF | 5.84E-15 |
| filter117⟷filter124 | SOX1⟷DMRTA2 | Sox⟷DM | 0.013489901 |
| filter109⟷filter162 | SOX1⟷DNMT1 | Sox⟷CxxC | 0.005835994 |
| filter73⟷filter161 | SOX1⟷EGR1 | Sox⟷C2H2 ZF | 0.003025464 |
| filter101⟷filter110 | SOX1⟷HHEX | Sox⟷Homeodomain | 0.003296044 |
| filter160⟷filter174 | SOX1⟷HOXA11 | Sox⟷Homeodomain | 3.04E-06 |
| filter159⟷filter160 | SOX1⟷KDM2B | Sox⟷CxxC | 0.012152077 |
| filter151⟷filter160 | SOX1⟷LCOR | Sox⟷Pipsqueak | 5.08E-08 |
| filter109⟷filter125 | SOX1⟷LHX5 | Sox⟷Homeodomain | 0.000158352 |
| filter117⟷filter131 | SOX1⟷SNAI3 | Sox⟷C2H2 ZF | 6.24E-05 |
| filter21⟷filter73 | SOX1⟷TLX2 | Sox⟷Homeodomain | 0.041759796 |
| filter117⟷filter195 | SOX1⟷ZNF263 | Sox⟷C2H2 ZF | 0.000640375 |
| filter160⟷filter192 | SOX1⟷ZNF35 | Sox⟷C2H2 ZF | 2.55E-05 |
| filter0⟷filter162 | SOX3⟷DNMT1 | Sox⟷CxxC | 0.040168612 |
| filter69⟷filter161 | SOX3⟷EGR1 | Sox⟷C2H2 ZF | 7.93E-05 |
| filter0⟷filter101 | SOX3⟷HHEX | Sox⟷Homeodomain | 1.05E-05 |

| | | | |
|---|---|---|---|
| filter1↔filter180 | SOX3↔HOXA6 | Sox↔Homeodomain | 0.017315501 |
| filter69↔filter160 | SOX3↔SOX1 | Sox↔Sox | 0.001305497 |
| filter21↔filter180 | SOX3↔TLX2 | Sox↔Homeodomain | 0.008802917 |
| filter69↔filter102 | SOX3↔ZNF263 | Sox↔C2H2 ZF | 4.12E-05 |
| filter148↔filter160 | TET1↔SOX1 | Sox↔CxxC | 8.61E-16 |
| filter21↔filter178 | TLX2↔ARID3B | Homeodomain↔ARID/BRIGHT | 0.042530051 |
| filter143↔filter193 | TLX2↔DMRTA2 | Homeodomain↔DM | 0.007554303 |
| filter162↔filter193 | TLX2↔DNMT1 | Homeodomain↔CxxC | 0.029581514 |
| filter48↔filter161 | TLX2↔EGR1 | Homeodomain↔C2H2 ZF | 0.006097183 |
| filter21↔filter92 | TLX2↔HMG20B | Sox↔Homeodomain | 0.02816662 |
| filter21↔filter174 | TLX2↔HOXA11 | Homeodomain↔Homeodomain | 0.001296995 |
| filter21↔filter199 | TLX2↔KDM2B | Homeodomain↔CxxC | 0.020374461 |
| filter151↔filter193 | TLX2↔LCOR | Homeodomain↔Pipsqueak | 7.29E-07 |
| filter21↔filter131 | TLX2↔SNAI3 | Homeodomain↔C2H2 ZF | 1.43E-05 |
| filter21↔filter148 | TLX2↔TET1 | Homeodomain↔CxxC | 3.12E-16 |
| filter21↔filter59 | TLX2↔ZKSCAN1 | Homeodomain↔C2H2 ZF | 0.019110844 |
| filter21↔filter57 | TLX2↔ZNF202 | Homeodomain↔C2H2 ZF | 0.002240944 |
| filter21↔filter195 | TLX2↔ZNF263 | Homeodomain↔C2H2 ZF | 0.0002947 |
| filter118↔filter193 | TLX2↔ZNF32 | Homeodomain↔C2H2 ZF | 0.004170142 |
| filter130↔filter192 | TLX2↔ZNF35 | Homeodomain↔C2H2 ZF | 6.62E-06 |
| filter59↔filter133 | ZKSCAN1↔DNMT1 | CxxC↔C2H2 ZF | 5.11E-16 |
| filter59↔filter101 | ZKSCAN1↔HHEX | Homeodomain↔C2H2 ZF | 3.90E-06 |
| filter59↔filter160 | ZKSCAN1↔SOX1 | Sox↔C2H2 ZF | 0.002379519 |
| filter57↔filter71 | ZNF202↔EGR1 | C2H2 ZF↔C2H2 ZF | 0.002000326 |
| filter57↔filter135 | ZNF202↔HHEX | Homeodomain↔C2H2 ZF | 0.021816047 |
| filter57↔filter174 | ZNF202↔HOXA11 | Homeodomain↔C2H2 ZF | 0.000574539 |
| filter57↔filter90 | ZNF202↔KDM2B | CxxC↔C2H2 ZF | 0.002906239 |
| filter57↔filter160 | ZNF202↔SOX1 | Sox↔C2H2 ZF | 5.26E-06 |
| filter57↔filter102 | ZNF202↔ZNF263 | C2H2 ZF↔C2H2 ZF | 0.000662677 |
| filter102↔filter107 | ZNF263↔ARID3B | C2H2 ZF↔ARID/BRIGHT | 1.63E-07 |
| filter174↔filter195 | ZNF263↔HOXA11 | Homeodomain↔C2H2 ZF | 3.53E-05 |
| filter102↔filter151 | ZNF263↔LCOR | C2H2 ZF↔Pipsqueak | 2.48E-11 |
| filter118↔filter133 | ZNF32↔DNMT1 | CxxC↔C2H2 ZF | 5.00E-18 |
| filter118↔filter138 | ZNF32↔KDM2B | CxxC↔C2H2 ZF | 5.45E-12 |
| filter118↔filter160 | ZNF32↔SOX1 | Sox↔C2H2 ZF | 0.003032589 |
